# SnoRNA Expression and RNA 2’-O-Methylation in *Drosophila melanogaster* S2 Cells

**DOI:** 10.64898/2026.05.21.726978

**Authors:** Xuan Ye, Yaling Liu, Sara Olson, Lijun Zhan, Gordon G. Carmichael, Brenton R. Graveley

## Abstract

Small nucleolar RNAs (snoRNAs) are a class of non-coding RNAs that play critical roles in guiding 2’-O-methylation (Nm) and pseudouridylation modifications of RNAs. In *Drosophila melanogaster*, snoRNAs undergo dynamic changes in expression during development. In this study, we identified 239 snoRNAs that are robustly expressed in *Drosophila* S2 cells, representing 87% of all annotated *Drosophila* snoRNAs. Given that box C/D snoRNAs guide site-specific 2’-O-methylation (Nm) of RNA, we next characterized the Nm landscape of S2 cells using RibOxi-seq2, a high-throughput approach capable of detecting Nm modifications with single-nucleotide resolution. RibOxi-seq2 revealed 17 Nm sites in 18S rRNA with a 94% concordance to previously reported RiboMeth-Seq data. In 28S rRNA, 30 Nm sites were identified, corresponding to an 71.4% overlap with established references. Additionally, we detected both a known Nm site (Gm74) and a novel site (Um66) in 5.8S rRNA, further validating the sensitivity and specificity of the approach. RibOxi-seq2 further identified Nm sites in small nuclear RNAs (snRNAs), expanding the annotation of modified non-coding RNAs. Additionally, the method revealed Nm modifications within internal regions of mRNAs. In total, we detected Nm modifications in 2,057 unique mRNAs, underscoring the widespread presence of this epitranscriptomic modification in coding transcripts. Strikingly, although we could not identify any snoRNAs predicted to guide the mRNA 2’-O-methylation modifications by canonical mechanisms, we identified strong consensus sequences surrounding many of these mRNA sites. Together, our findings not only expand the known landscape of Nm-modified RNAs but also highlight the robustness of RibOxi-seq2 for transcriptome-wide RNA modification profiling. Collectively, this study presents a comprehensive atlas of snoRNA expression and 2’-O-methylation sites in *Drosophila* S2 cells, offering valuable insights into the epitranscriptomic landscape orchestrated by snoRNAs.

## Introduction

Small nucleolar RNAs (snoRNAs) represent a highly conserved and abundant family of functional non-coding RNAs, typically ranging from 49 to 511 nucleotides in length (1–3). In higher eukaryotes, including *Drosophila* and humans, the vast majority of snoRNAs (>90%) are transcribed, excised, and processed from the intronic regions of protein-coding or long non-coding host genes (1–3).

Based on their conserved structural motifs and interactions with specific core proteins to form functional small nucleolar ribonucleoprotein (snoRNP) complexes, snoRNAs are classically divided into two primary families: the box C/D and the box H/ACA snoRNAs (**Figure 1A**) (3, 4). Box C/D snoRNAs consist of conserved C/C’ and D/D’ sequence motifs that fold into a kink-turn structure, facilitating the recruitment of core proteins such as NOP56, NOP58, SNU1, and the methyltransferase fibrillarin (FBL). The box C/D snoRNAs primarily guide the site-specific 2’-O-methylation of RNA exactly five nucleotides upstream of the D or D’ box (2, 5, 6). Conversely, box H/ACA snoRNAs form a characteristic hairpin-hinge-hairpin-tail structure and assemble with the pseudouridine synthase dyskerin, as well as NHP2, NOP10, and GAR1, to catalyze the conversion of the targeted uridines into pseudouridines (6). These targeted modification events are critical for the structural stability, pre-rRNA folding, and functional optimization of cellular RNAs, ultimately impacting translational fidelity and ribosome biogenesis. In addition to canonical families, a specialized subset known as small Cajal body RNAs (scaRNAs), localize to nuclear cajal bodies and are featured with discrete localization signals such as the CAB box or GU/UG wobble stems. scaRNAs are responsible for guiding the pseudouridylation and 2’-O-methylation of spliceosomal small nuclear RNAs (snRNAs), thereby playing a critical role in spliceosome maturation and function (6, 7).

**Figure 1.**
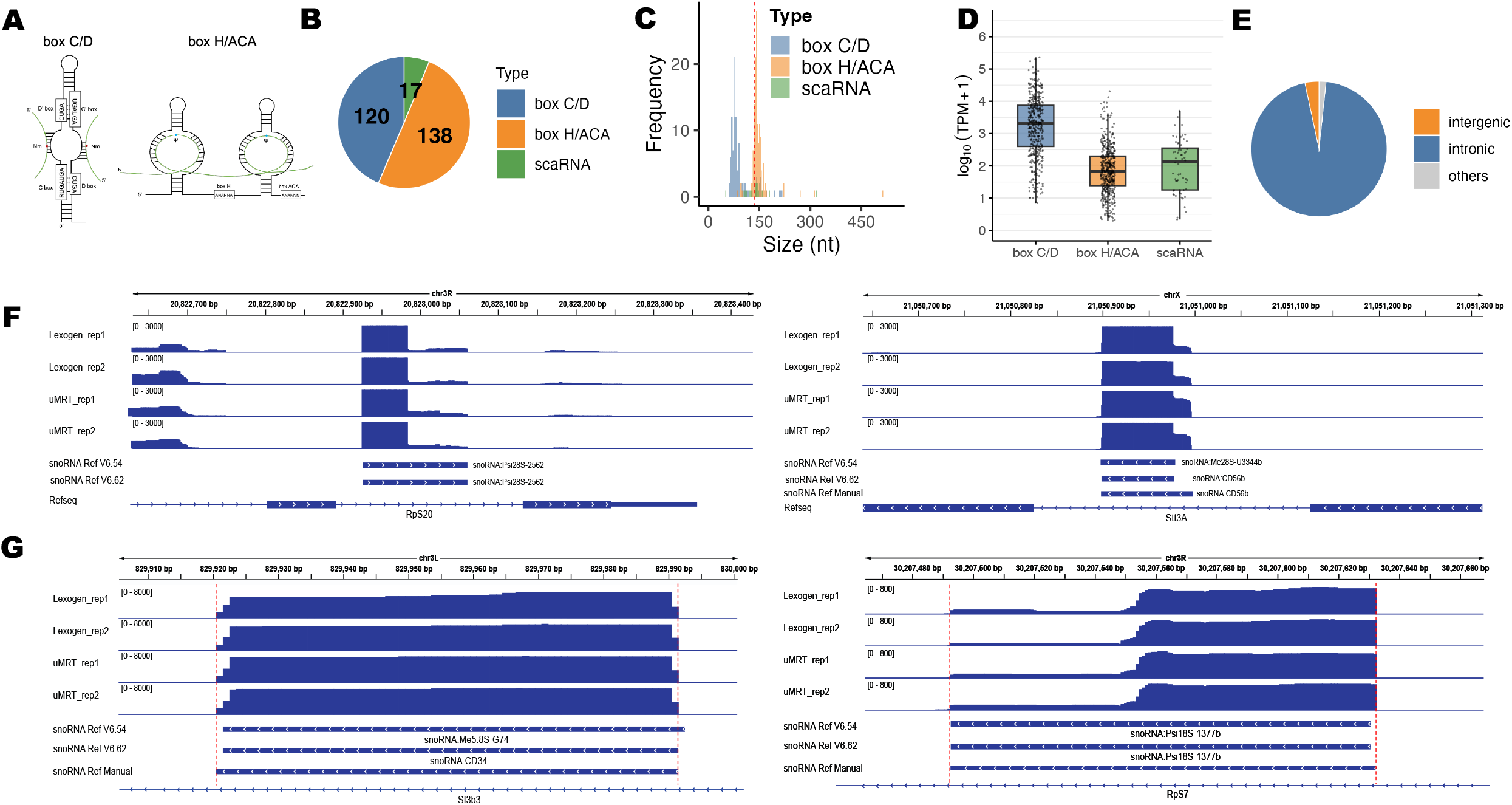
SnoRNA Landscape in *Drosophila* S2 Cells. A. Schematic of box C/D and box H/ACA snoRNA. B. Distribution of snoRNAs classes annotated in the *Drosophila* genome reference. C. Length distribution of snoRNAs, color-coded by class (box C/D, box H/ACA, scaRNA). D. Heatmap of the snoRNA expression level [log2(TPM+1)] for 4 technical replicates, rows are grouped by the snoRNA class. E.The genomic context of the snoRNAs, color-coded by type. F. Representative snoRNA expression profiles from small RNA-Seq data. Multiple genome annotation versions are shown, snoRNA nomenclature differs between annotation releases. G. Representative snoRNA expression profiles from small RNA-Seq data. Multiple genome annotation versions are shown, snoRNA nomenclature differs between annotation releases. Manually curated snoRNA boundaries are indicated by red dashed lines.

Beyond their canonical roles in rRNA and snRNA modifications, the functional landscape of snoRNAs has expanded significantly in recent years. It is now recognized that snoRNAs can also guide modifications on tRNAs and mRNAs, and even participate in rRNA acetylation (8). Furthermore, a large proportion of snoRNAs lack identified rRNA or snRNA targets and are termed “orphan” snoRNAs (1, 3). These, along with other canonical snoRNAs, exhibit diverse non-canonical regulatory functions, including the regulation of pre-mRNA alternative splicing, modulation of mRNA 3’ end processing and stability, and the maintenance of chromatin accessibility (9). Mature snoRNAs can also be processed into shorter, stable snoRNA-derived RNAs (sdRNAs) that display microRNA-like or piRNA-like regulatory activities to influence gene expression (10). Consequently, snoRNA dysregulation is heavily implicated in cellular stress responses, such as lipotoxicity, cancers and Prader-Willi syndrome (11–13). Importantly, snoRNA expression undergoes dynamic changes during development and differentiation. In *Drosophila melanogaster*, for example, snoRNA expression patterns vary across developmental stages, highlighting their functional plasticity and potential contributions to organismal growth (14).

2’-O-methylation (Nm) is among the most prevalent RNA modifications in eukaryotes (15). Nm involves the addition of a methyl group to the 2’ hydroxyl position of the ribose sugar, conferring resistance to nuclease degradation, enhanced RNA folding and thermostability, and altered RNA-protein interactions. Nm is particularly enriched in rRNA, where it plays essential roles in ribosome biogenesis and translational fidelity (9). Beyond rRNA, Nm modifications have been identified on snRNAs and tRNAs, and more recently on mRNAs, where they are proposed to modulate transcript stability and translation efficiency (16–19). The deposition of Nm marks is primarily guided by box C/D snoRNAs, which base-pair with target RNAs and recruit the methyltransferase FBL to catalyze methylation at a precisely defined nucleotide position (20).

A diverse repertoire of high-throughput sequencing strategies has been developed to systematically map Nm, each capable of single-nucleotide resolution but differing in chemistry, scope, and sensitivity (9). RiboMeth-seq, the earliest of these methods, exploits the resistance of Nm residues to alkaline hydrolysis to generate quantitative methylation profiles and remains uniquely suited for robust stoichiometric measurement of rRNA and other highly abundant non-coding RNAs (21). 2’-OMe-seq instead leverages the propensity of reverse transcriptase (RT) to stall at Nm residues under low-dNTP conditions, enabling systematic detection of Nm signatures, although sensitivity is likewise biased toward abundant RNA species (22). Nm-seq and RibOxi-seq2 couple RNA fragmentation with sodium periodate oxidation and β-elimination to selectively degrade fragments with unmodified 3’ RNA termini, thereby enriching Nm-protected fragments for sequencing (16, 17, 23–25). More recently, Nm-mut-seq has employed engineered HIV RT variants that traverse Nm sites with elevated dATP misincorporation, enabling transcriptome-wide mapping at the cost of reduced sensitivity at certain modifications such as Um (18). Complementing these RT- and chemistry-based approaches, NJU-seq uses Mycoplasma genitalium RNase R, which degrades RNA in the 3’ to 5’ direction but stalls precisely one nucleotide downstream of Nm, producing highly specific cleavage signatures. Its broader adoption, however, has been constrained by an elevated false-positive rate and the lack of a commercial enzyme source (26). In parallel, Nanopore direct RNA sequencing (DRS) has emerged as a promising orthogonal strategy for detecting Nm on native transcripts, particularly when pairing wild-type and Nm-deficient samples, and offers the additional advantages of long-read, isoform-resolved profiling (27–30).

Transcriptome-wide Nm profiling has been concentrated in mammalian and yeast systems, leaving the Nm landscape of many important model organisms, including *Drosophila* melanogaster, largely uncharted. In this study, we selected RibOxi-seq2 to map Nm modifications across the *Drosophila* transcriptome, given its single-nucleotide resolution and broad coverage across diverse RNA classes. To biochemically validate sites identified by RibOxi-seq2, we employed the quantitative tool Nm-VAQ (Nm Validation and Absolute Quantification), which leverages the ability of 2’-O-methylation to impede RNase H cleavage of RNA-DNA hybrid substrates (26). In this assay, a chimeric RNA-DNA probe directs RNase H to a specific position within the target RNA, and the methylation-dependent reduction in cleavage efficiency is quantified by qPCR, yielding absolute Nm stoichiometry at the queried site. Together, these complementary strategies enabled a comprehensive mapping of Nm modifications across the *Drosophila* transcriptome and the integration of these sites with their candidate snoRNA guides.

Here we combined small RNA sequencing with RibOxi-seq2 to comprehensively profile snoRNA expression in a *Drosophila* cell line, S2. This integrative strategy enabled us to generate a global atlas of snoRNA abundance and Nm modification profiles, uncovering both evolutionarily conserved and previously unreported sites. Collectively, our results provide a robust framework for Nm site annotation in *Drosophila* and establish a valuable resource for advancing snoRNA biology, exploring the epitranscriptomic landscape, and facilitating comparative studies across species.

## Results

### SnoRNA Expression in *Drosophila* S2 Cells

To establish a robust reference for our snoRNA analysis, annotations were retrieved from the *D. melanogaster* FlyBase database, the most recent at the time of this study being dmel_r6.67 (31). We note that a major nomenclature change was introduced in 2024 with release dmel_r6.58 (FB2024_03). In dmel_r6.58, all 120 box C/D snoRNAs were systematically renamed to resemble human nomenclature. For example, previously annotated snoRNAs in dmel_r6.54 such as snoRNA:Me28S-G1083c and snoRNA:Or-CD9a were renamed to snoRNA:CD28c and snoRNA:CD63a in dmel_r6.58, respectively (27). Furthermore, the genomic coordinates for 93 snoRNAs were refined to provide more accurate transcript boundaries (**Table S1**). Despite these improvements, a duplicated genomic record for scaRNA:PsiU2-55 persists in the genome reference (**Table S1**), highlighting the need for continued curation of the *Drosophila* snoRNA reference.

The dmel_r6.67 reference contains a total of 275 annotated snoRNA species, including 120 box C/D snoRNAs, 138 box H/ACA snoRNAs, and 17 scaRNAs (**Figure 1B**). The transcript lengths of these snoRNAs range from 46 to 511 nt, with an average length of approximately 120 nt (**Figure 1C**). One notable outlier, snoRNA:ACA9, is annotated at an unusual length of 511 nt, which is 195 nt longer than scaRNA:MeU5-C46, the second longest annotated snoRNA. Overall, the box H/ACA snoRNAs averaged 143 nt in length, approximately 52 nt longer than box C/D snoRNAs (mean 91 nt).

To generate a comprehensive atlas of snoRNA expression in *Drosophila* S2 cells, we profiled the small RNA transcriptome in four independent technical replicates. Libraries were constructed with the Lexogen Small RNA-Seq protocol, which employs two successive rounds of adapter ligation to the 3’ and 5’ ends of target RNAs, followed by reverse transcription and amplification for library construction. Because snoRNAs adopt extensive secondary structures that can hinder reverse transcription, we evaluated two RT enzymes in parallel, two replicates were prepared using the standard Lexogen reverse transcriptase, while the other two used UltraMarathon (uMRT), a highly processive Group II intron reverse transcriptase which can efficiently resolve structural templates and read through certain RNA modifications (32) (**Figure S1A, S1B**). We hypothesized that utilizing uMRT would minimize premature termination during reverse transcription, thereby enabling a more complete and accurate representation of the snoRNA transcriptome.

Both the Lexogen and the uMRT enzyme protocol successfully captured the diversity of small RNA species expressed in *Drosophila* S2 cells, including snoRNAs, tRNAs, rRNA fragments, and other ncRNAs (**Figure S1C**). While uMRT showed clear advantages for tRNA quantification, consistent with its enhanced processivity through complex secondary structures (**Figure S1D**), both uMRT and the Lexogen kit performed comparably for snoRNA characterization, with a correlation coefficient of 0.91 (**Figure S1E**). We detected 246 snoRNAs expressed at a TPM > 1, corresponding to approximately 89% of all snoRNAs annotated in the current reference (dmel_r6.67) (**Figure 1D**). The remaining 11% are most likely not highly expressed in S2 cells rather than missed by our protocol, as snoRNA expression is known to be dynamically regulated across cell types and developmental stages (33). Consistent with two recent studies using TGIRT that characterized *Drosophila* snoRNA expression (2, 27), no novel snoRNAs were detected, supporting the completeness of the current annotation. Additionally, snoRNAs that were present in dmel_r6.54 but were subsequently removed from dmel_r6.58 were also not detected in our data, supporting the reliability of the current genome annotations.

As has been previously reported (34), an analysis of the genomic organization of the expressed snoRNAs revealed that 95% of them are encoded within the introns of host genes, while only ∼3% are located in intergenic regions (**Figure 1E**). Notably, many of the intronic snoRNAs are organized into well-characterized clusters, such as those within the *Uhg1*–*Uhg8, lola, Arpc2* and *dom* gene loci (**Table S1, Figure S1F**). Moreover, 30% (73 of the 239) of the snoRNA host genes encode ribosomal proteins, such as RpL14, RpL17, RpS5a, and RpS7 (**Table S1**).

The ligation-based small RNA sequencing strategy also enabled the detection of snoRNA processing intermediates, providing insights into snoRNA maturation pathways. For example, in snoRNA:Psi28S-2562, sequencing profiles consistently revealed read populations that are substantially shorter than the annotated reference region. The 58 nt fragment emerged as a stably expressed snoRNA-derived fragment, suggesting that it may represent a processed and functionally relevant species rather than simple degradation product. In contrast, for snoRNA:Me28S-U3344b, the coverage tracks revealed a subset of reads that extended well beyond the annotated reference (**Figure 1F**). These longer transcripts may represent precursor molecules of the snoRNA, suggesting intermediate forms in the maturation pathway and providing insights into the processing steps of snoRNA biogenesis.

End-to-end read coverage further enabled refinement of annotated snoRNA loci. The 5′ start site of *snoRNA:Me5*.*8S-G74* (*snoRNA:CD34*) was corrected between FlyBase GTF versions 6.58 and 6.62, and we manually extended its 3′ end based on our data.

Likewise, the 5′ start site of *snoRNA:Psi18S-1377b* was revised based on small RNA-seq read coverage (**Figure 1G**). Overall, by systematically comparing sequencing coverage with FlyBase GTF records, we manually curated and corrected discrepancies in the transcript boundaries of 174 snoRNAs, representing 72% of highly expressed snoRNAs (**Table S2**). The revised annotation was supported by concordant read coverage across libraries prepared with both the Lexogen RT and uMRT. Together, these findings highlight the importance of integrating experimental evidence to further refine snoRNA atlas, ensuring accurate representation of their true transcript structures.

### Landscape of Nm sites on rRNA

To generate an Nm map of the *Drosophila* transcriptome, we performed RibOxi-seq2 (**Figure 2A**) using mRNA-enriched from *Drosophila* S2 cells. We found that the residual amount of rRNA remaining in these samples provided sufficient coverage to concurrently profile rRNA modifications, which provided an important quality control for RibOxi-seq2. Importantly, because RibOxi-seq2 is qualitative rather than quantitative, we could not determine the stoichiometry of the Nm levels at each position from RibOxi-Seq2 alone. To define confident Nm peaks in the data, we therefore applied an empirical cutoff of 500 reads per position.

**Figure 2.**
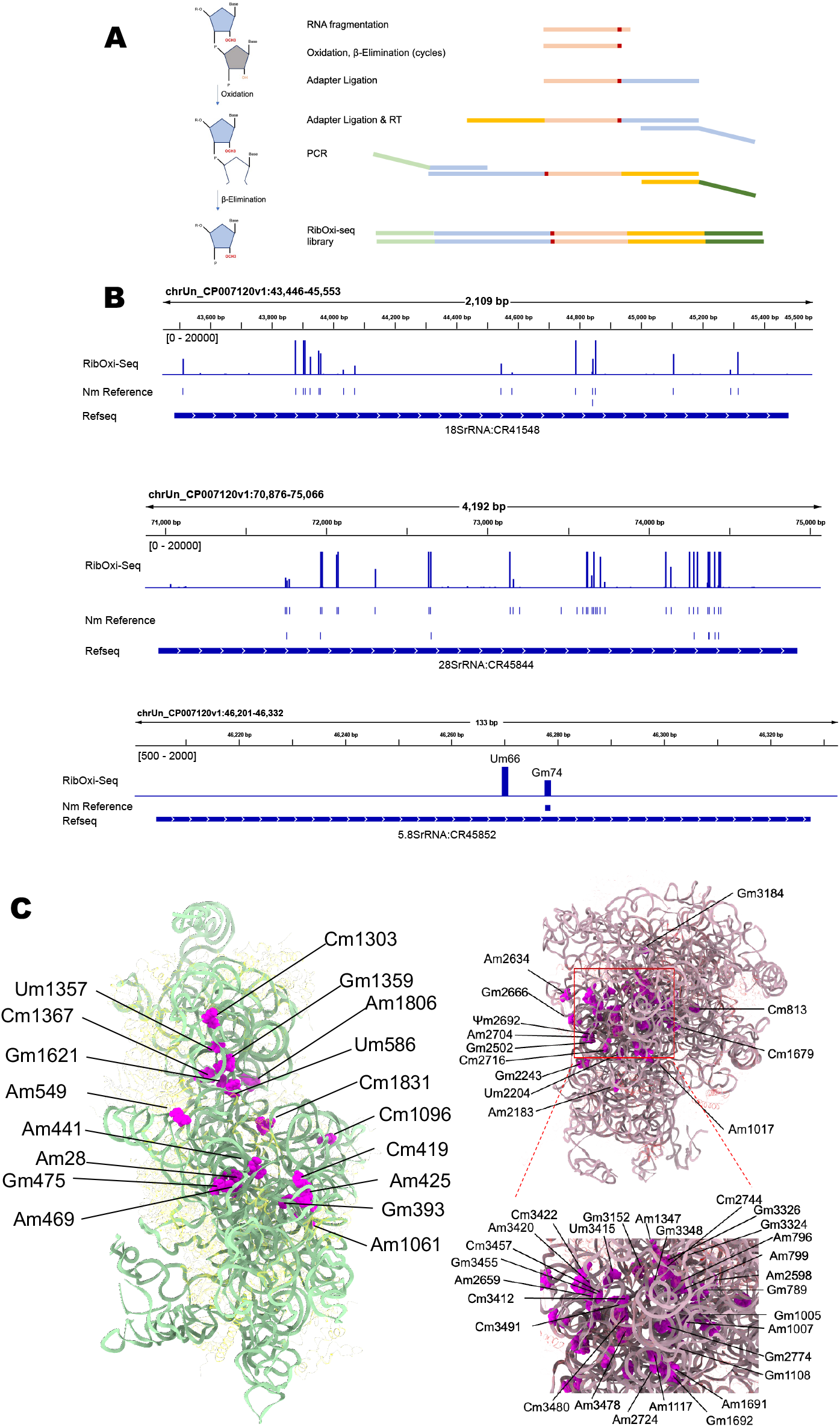
RibOxi-Seq2 detection of 2’-O-methylation in rRNAs in *Drosophila* S2 cells. A. Chemistry of NaIO_4_ treatment of RNAs bearing a free 2’-OH group. The oxidation step opens the ribose ring, and the subsequent β-elimination step removes the terminal nucleotide, exposing RNAs with a 2’-O-methylated 3’-end that are resistant to oxidation. NaIO_4_ RNAs are then ligated to sequencing adapters and processed through the RibOxi-seq2 library preparation pipeline (see Methods for details). B. RibOxi-Seq2 peaks in 18S, 28S and 5.8S rRNA. Each peak aligns with validated or annotated Nm reference. C. Nm sites identified by RibOxi-seq2 mapped onto the cryo-EM (PDB:6XU6) structure of the *Drosophila* 60S and 80S ribosomal subunit. Nm-modified nucleotides are shown as magenta spheres; the 18S rRNA is shown as a green ribbon and ribosomal proteins as yellow ribbons; the 28S rRNA is shown as a purple ribbon and ribosomal proteins as red ribbons.

The *Drosophila* rRNA Nm landscape has been previously curated in the Modomics databases (35), and more recently expanded through integration of RiboMeth-Seq with Nanopore direct RNA sequencing (DRS), which produced a comprehensive atlas of rRNA Nm modifications across S2R+ cells and wild-type Canton-S tissues (27). We leveraged these resources as an independent benchmark for the performance of RibOxi-seq2.

In 18S rRNA, RibOxi-seq2 detected 17 of the 18 previously annotated Nm sites (94% recovery), demonstrating the nucleotide-level precision of this approach (**Figure 2B, Figure S2A, Table S3**) (27, 35). In 28S rRNA, 30 of 42 annotated sites were recovered (71.4% overlap), along with one previously unreported modification at Um814 (**Figure 2B, Figure S2B**). The slightly lower concordance in 28S is attributable to a known limitation of periodate-based methods at closely spaced Nm positions. For example, at positions Gm3324/Gm3326, oxidative cleavage is arrested at the first 2’-O-methylated nucleotide, leaving the immediately adjacent site undetected. Mapping detected sites onto the *Drosophila* 80S ribosome cryo-EM structure (PDB:6XU6) revealed evident spatial clustering within functionally critical regions, including the peptidyl transferase center (**Figure 2C**) (36). In 5.8S rRNA, we confirmed the known Gm74 site and identified a novel Um66 modification not annotated in prior reference datasets (**Figure 2B**) (27). In the mitochondrial large subunit (mt-LSU) rRNA, three Nm sites (Um245, Am433, and Gm1179) were detected (**Figure S2B**), of which Gm1179 coincides with a position previously reported by Nanopore-based profiling (27). Together, these observations across cytoplasmic and mitochondrial rRNAs demonstrate both the sensitivity and single-nucleotide precision of RibOxi-seq2 for comprehensive Nm mapping.

To biochemically validate the RibOxi-seq2 peaks we observed in rRNA, we selected representative Nm sites spanning the *Drosophila* rRNAs and quantified their modification levels using Nm-VAQ (**Figure S3A**). As a positive control, we first assayed the well-characterized Cm1367 site in 18S rRNA and observed a modification level of approximately 95%, confirming both the assay’s quantitative accuracy as it is consistent with the near-complete methylation of this reference site. Notably, the orthologous position in human 18S rRNA is not 2’-O-methylated but instead carries an N4-acetylcytidine (ac4C) modification (37). Because ac4C acetylates the exocyclic N4 amine of the cytosine base and leaves the ribose 2’-OH chemically unchanged, it is not detected by RibOxi-seq2. The distinct modification identities at this evolutionarily conserved position therefore reflects a biological divergence between *Drosophila* and humans, implying that nucleotide-level sequence conservation does not necessarily predict conservation of the underlying chemical modification.

We next applied Nm-VAQ to the newly identified Um66 site in 5.8S rRNA, and observed a modification frequency of approximately 76%, indicating near-constitutive deposition at this position (**Figure S3B**). Multiple sequence alignment across a diverse set of eukaryotes revealed complete conservation of the U66 residue, consistent with an essential functional role and suggesting that 2’-O-methylation at this position is likely preserved in other eukaryotic species.

Following the annotation and validation of the rRNA Nm sites, the canonical snoRNAs predicted to catalyze these modifications were identified using Snoscan2 (38). Table S1 lists a mapping of each Nm site to its corresponding snoRNA was subsequently generated. Notably, we found several discrepancies relative to previous predictions (27), and a subset of rRNA Nm sites could not be assigned to any candidate snoRNA. Furthermore, our analysis suggests that snoRNA isoforms sharing high sequence similarity do not necessarily guide the same Nm modifications. For instance, snoRNA:CD5b and snoRNA:CD5c, but not snoRNA:CD5a, were predicted to guide Am28 methylation on 18S rRNA, highlighting isoform-specific functional divergence within the same snoRNA family.

To assess the evolutionary architecture of rRNA 2’-O-methylation, we compared the *Drosophila* Nm landscape with published maps in human rRNAs (**Figure S2C**). While many Nm sites are conserved, distinct species-specific modification regions were observed. For instance, human 28S rRNA harbors specific Nm signals within regions 398–400, 2401–2804, and 4571–4731, whereas *Drosophila* 28S rRNA exhibits a fly-specific Nm cluster at positions 3415–3420 (**Figure S2D**). Similarly, human 18S rRNA contains unique Nm peaks in regions 99–436, 644–867, and 1383–1668, further illustrating the evolutionary divergence of rRNA modification profiles. These findings reveal a mosaic of highly conserved and species-specific 2’-O-methylation patterns across *Drosophila* and human rRNAs.

### Landscape of Nm sites on other ncRNAs

While RiboMeth-Seq is highly effective for profiling Nm modifications in abundant non-coding RNAs (such as rRNAs), it generally lacks the sensitivity to detect modifications on low-abundance transcripts at standard sequencing depths. In contrast, RibOxi-seq2 enables broader transcriptomic coverage. Applying an empirical peak threshold of ≥500 reads, we identified Nm modifications distributed across snRNAs, tRNAs, lncRNAs, and mRNAs in addition to the canonical rRNA substrates, highlighting the pervasiveness of ribose methylation across the *Drosophila* transcriptome (**Figure 3A**).

**Figure 3.**
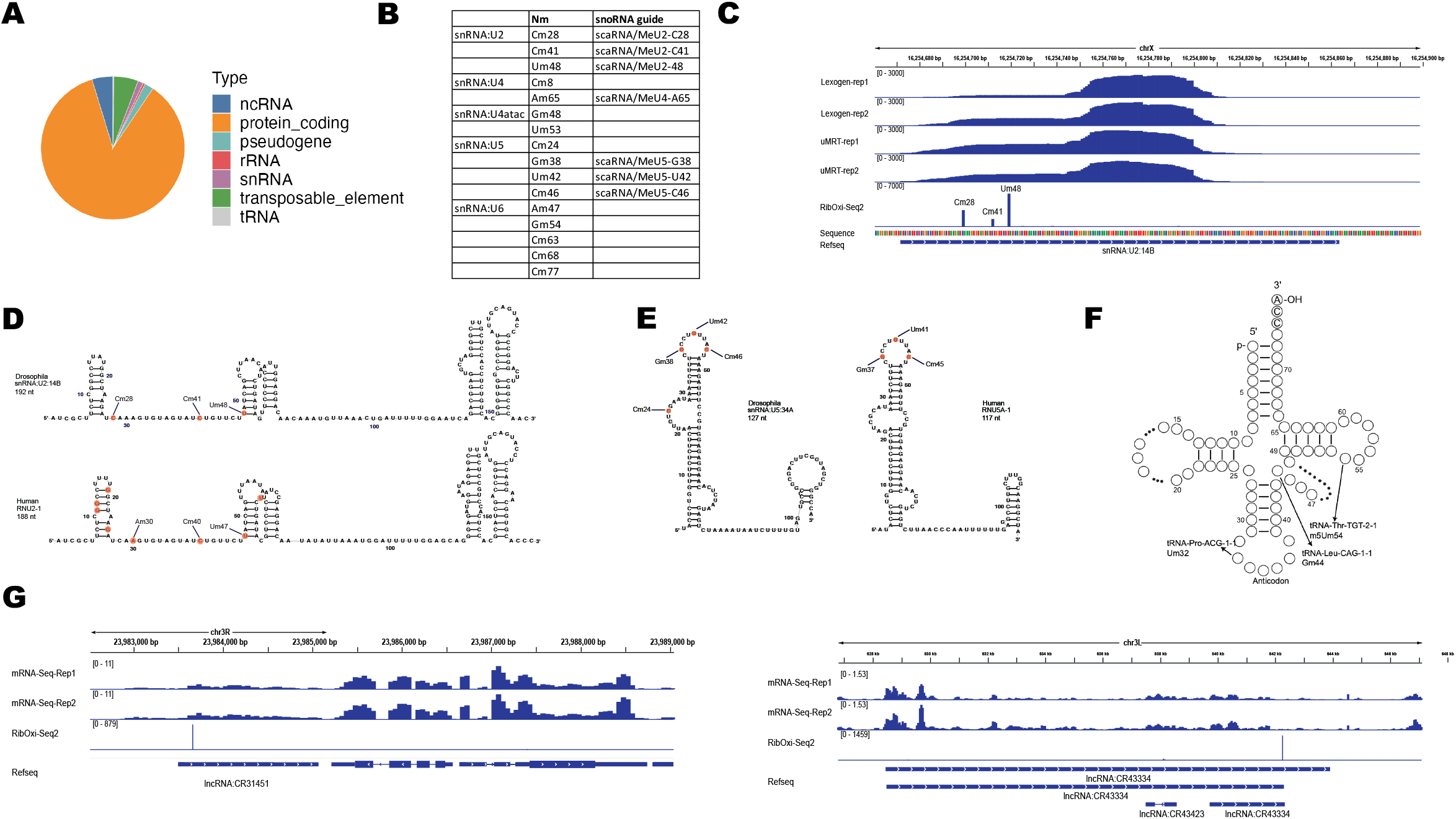
RibOxi-Seq2 detection of 2’-O-methylation in snRNAs, tRNAs and lncRNAs in *Drosophila* S2 cells. A. Pie chart showing the distribution of Nm sites across RNA biotypes. B. Summary of Nm sites identified in *Drosophila* snRNAs by RibOxi-seq2, with their corresponding guide snoRNAs. C. Representative view of U2 snRNA showing small RNA-seq expression and RibOxi-seq2 signal. Each RibOxi-seq2 peak corresponds to a previously validated or annotated Nm modification site. D. Comparison of U2 snRNA secondary structures from *Drosophila* and human. Nm sites identified in *Drosophila* by RibOxi-seq2 are highlighted in orange; human Nm sites were obtained from prior annotations. E. Comparison of U5 snRNA secondary structures from *Drosophila* and human. Nm sites identified in *Drosophila* by RibOxi-seq2 are highlighted in orange; human Nm sites were obtained from prior annotations. F. Summary of Nm sites identified in *Drosophila* tRNA by RibOxi-seq2, including the m5Um modification. G. Representative view of U2 snRNA showing small RNA-seq expression and RibOxi-seq2 signal.

A diverse landscape of Nm modifications across multiple snRNA species in *Drosophila* were mapped (**Figure 3B**). In U2 snRNA, three Nm sites were detected by RibOxi-Seq2: Um48, Cm41, and Cm28 (**Figure 3C**). These three sites were independently validated by Nm-VAQ which revealed them to be modified at near-constitutive levels with Um48 at 76%, Cm41 at 94%, and Cm28 at 100%, all of which are consistent with previously characterized sites in other eukaryotes (39–41). Notably, Gm25 and Am31 were not detected, as RibOxi-seq2 can not differentiate closely spaced Nm sites. snRNA Nm deposition is generally guided by scaRNAs, and the scaRNAs predicted to guide modification at each validated site are listed in Figure 3B. Comparative analysis of the predicted *Drosophila* and human U2 snRNA secondary structures revealed that while *Drosophila* Cm28 and Cm41 occupy divergent sequence contexts relative to their human counterparts, *Drosophila* Um48 is positionally conserved between the two species, mapping to the first nucleotide of the second stem-loop (**Figure 3D**).

Comparison against the known human snRNA modification repertoire revealed both evolutionarily conserved and lineage-specific Nm features. In *Drosophila* U5 snRNA, three conserved Nm sites (Gm38, Um42, and Cm46) were mapped within the stem-loop region, each aligned to its human ortholog (Gm37, Um41, and Cm45) (**Figure 3E, S4A**). Strikingly, we also identified a previously unreported Cm modification at position 24 (Cm24), located within the homologous stem-loop, which is 3 nt longer in *Drosophila* than in human (**Figure 3E, S4A**). Notably, the *Drosophila* ortholog of this stem-loop is 2 nt larger compared to its human counterpart, suggesting that Cm24 is a lineage-specific modification that may contribute to the structural or functional fine-tuning of U5 snRNP assembly or splicing in *Drosophila*.

Nm modifications were also identified across multiple tRNA species. To account for variable arm size differences among tRNA isotypes, nucleotide positions were assigned according to the universal tRNA numbering system (42). Canonical Nm sites were detected at position 32 in multiple tRNAs, including tRNA-Pro-AGG-1-1 and tRNA-Pro-CGG-1-1 (**Figure 3F, Table S3**). Modification at this position is catalyzed by CG5220, a homolog of yeast TRM7 and human FTSJ1, consistent with the evolutionary conservation of FTSJ-family-mediated 2’-O-methylation across eukaryotes. Additionally, Gm44 modification was detected in tRNA-Leu-CAG-1-1, tRNA-Leu-CAG-1-2 (**Figure 3F, Table S3**). This 2’-O-methylation is linked to CG9386, the *Drosophila* homolog of yeast Trm44. The modification m5Um (5,2’-O-dimethyluridine) was mapped at position 54 in tRNA-Thr-TGT-2-1 (**Figure 3F, Table S3**), matching our prior human transcriptome-wide findings (17, 43). As expected, sodium periodate treatment cannot distinguish Nm from m5Um, as the latter represents a ribose modification coupled to a base modification. Lastly, due to the 18-nucleotide minimum read length required for mapping in RibOxi-seq2, our approach inherently cannot map closely clustered proximal sites. Consequently, we did not detect the common Nm sites at positions 18, 20, or 34.

Notably, RibOxi-seq2 also detected 2’-O-methylation events on lncRNA species, including lncRNA:CR31451 and lncRNA:CR43334, representing a previously uncharacterized layer of lncRNA modification in *Drosophila* (**Figure 3G)**. Notably, both lncRNAs have well-characterized biological roles. lncRNA:CR31451 functions as a neuronal architectural RNA scaffolding Staufen-containing granules required for synaptic signaling and nervous system maturity (44), while lncRNA:CR43334 is the precursor transcript of the bantam miRNA, a central regulator of cell proliferation, apoptosis, and tissue growth in *Drosophila* (45). The detection of Nm modifications on these lncRNAs raises the possibility that 2’-O-methylation may contribute to their processing, stability, or regulatory activity.

Together, these results demonstrate that RibOxi-seq2 enables consistent and high-resolution detection of Nm sites across diverse RNA classes. The data are consistent with existing annotations, and, when combined with the identification of previously unreported sites and RNA types, support both the accuracy of the approach and its utility as a comprehensive tool for transcriptome-wide Nm profiling.

### Landscape of Nm sites on mRNAs

The high sensitivity of RibOxi-seq2 has previously been performed to map Nm modifications within internal regions of mRNAs (17, 23, 24). To characterize the mRNA 2’-O-methylation landscape in *Drosophila* S2 cells, we first enriched the poly(A) mRNA fraction and performed parallel RNA-seq to obtain transcript-level expression profiles. RibOxi-seq2 reads mapping to mRNA regions were then normalized to transcript expression levels (TPM) to account for abundance differences, and mRNA Nm sites were defined using a signal-to-expression ratio cutoff of 2. Applying this threshold, we identified 2,057 mRNAs bearing prominent Nm signals (**Table S3**), demonstrating that 2’-O-methylation is pervasive across the mRNA transcriptome. The majority of methylated genes contained only a single Nm site (73.9%) and 18.8% contained two sites, the remaining 7.3% harbored three or more sites; one outlier gene, *Ubi-p63E*, contained as many as 20 Nm sites (**Figure 4A**). The number of mRNA Nm sites per gene identified is comparable to that previously reported in human HeLa cells (16).

**Figure 4.**
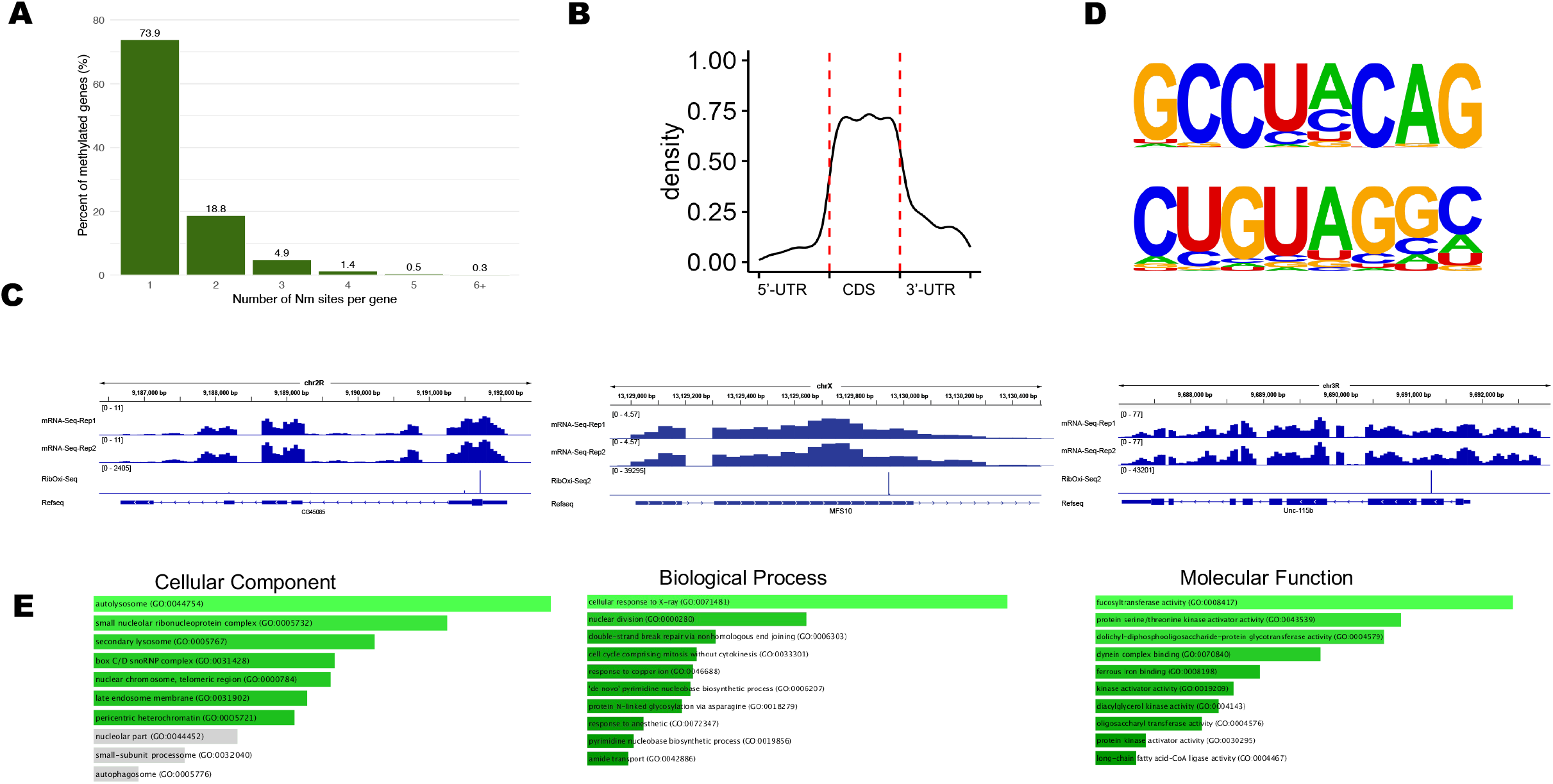
RibOxi-Seq2 detection of 2’-O-methylation in mRNAs in *Drosophila* S2 cells. A.The percentage of methylated genes according to the number of Nm sites per gene. B. Metagene profile of Nm sites distribution along a mRNA transcript. C. Representative view of mRNA showing small RNA-seq expression and RibOxi-seq2 signal. D.Top-ranked sequence motif identified by HOMER de novo motif analysis of the 500 highest-confidence *Drosophila* Nm sites. The motif logo and associated p-value are shown. E. Gene Ontology enrichment analysis of *Drosophila* mRNAs harboring Nm modifications.

Gene body analysis revealed that Nm sites were predominantly localized within the coding sequence (CDS), with weaker enrichment in the 3′ UTR, and the weakest enrichment in the 5’ UTR (**Figure 4B**). This distribution pattern closely mirrors prior transcriptome-wide characterizations of mRNA Nm in human embryonic stem cells and neurons as well as other cultured cells (16, 17), suggesting that preferential enrichment of 2’-O-methylation within coding regions may represent an evolutionarily conserved feature of the mRNA epitranscriptome.

To validate the RibOxi-seq2 identified mRNA Nm sites, we applied Nm-VAQ to three candidate transcripts, *mfs10* (a predicted transmembrane transporter involved in phosphate ion transport), *unc-115b* (an actin-binding protein with roles in lamellipodium assembly), and *CG45085* (an uncharacterized protein-coding gene) (**Figure 4C, Figure S3B**). Nm modifications at all three sites were successfully confirmed, with stoichiometries ranging from 34% to 72%. The considerable variation in Nm stoichiometry across transcripts indicates that mRNA 2’-O-methylation is not uniformly deposited and suggests that individual modification events may be dynamically regulated in a transcript-specific manner, consistent with recent reports of sub-stoichiometric mRNA modifications in other systems (17, 19).

The canonical pathway for Nm deposition on rRNA and snRNA relies on box C/D snoRNP and scaRNP complexes, in which a snoRNA guide base-pairs with the target RNA to position FBL for methylation. To test whether mRNA Nm sites are similarly installed through this pathway, we computationally screened the annotated *Drosophila* snoRNA pool against each mRNA Nm site for base-pairing complementarity compatible with canonical box C/D snoRNA–target interactions. Surprisingly, no significant complementarity could be established for any of the identified mRNA Nm sites, indicating that mRNA 2’-O-methylation in *Drosophila* S2 cells may not proceed through conventional snoRNA-directed targeting. These results point to the possible involvement of alternative mechanisms or enzymatic activities capable of specifically recognizing mRNA substrates, highlighting a potentially distinct and snoRNA-independent layer of 2’-O-methylation regulation in coding transcripts.

To further investigate the sequence context of these sites, we performed motif analysis on the top 500 Nm sites using HOMER. A significant sequence motif, CUGUAGGC (p = 1×10^-68^), was identified in 24.4% of target sequences compared to only 3.17% of background sequences (**Figure 4D**). However, neither this motif nor its reverse complement shares meaningful sequence similarity with the consensus box C/D snoRNA C box motif (RUGAUGA), further suggesting that Nm deposition in mRNA may not be guided by canonical snoRNA-directed mechanisms. Additionally, searches against the ATtRACT RNA-binding protein motif database also failed to identify any annotated RBP recognition motif matching this sequence, suggesting this sequence motif is targeted by an uncharacterized RBP or a guide-independent methyltransferase (46).

To investigate the functional relevance of 2’-O-methylation on mRNAs, we performed Gene Ontology (GO) enrichment analysis on the top 40 mRNAs with the highest 2’-O-methylation signals using FlyEnrichr (**Figure 4E**). Cellular Component analysis revealed significant enrichment for snoRNP-related complexes, including the box C/D snoRNP complex, small nucleolar ribonucleoprotein complex, and nucleolar components, suggesting that mRNAs encoding the 2’-O-methylation machinery itself are preferentially modified. Biological Process enrichment highlighted DNA damage response and cell cycle regulation, with top terms including cellular response to X-ray irradiation, double-strand break repair via non-homologous end joining, and nuclear division. Molecular Function analysis identified a coherent cluster of glycosylation-related enzymes, including fucosyltransferase, oligosaccharyltransferase, and dolichyl-diphosphooligosaccharide-protein glycotransferase activities. Together, these findings suggest that box C/D snoRNA-guided 2’-O-methylation preferentially targets mRNAs involved in snoRNP biogenesis, DNA damage response, and glycosylation, pointing to potential regulatory roles of this modification in maintaining cellular homeostasis.

To investigate the evolutionary conservation of mRNA Nm between human and *Drosophila* species, we examined several specific genes. *TOP1* and *ACTG1* were previously reported to be internally 2’-O-methylated in human CT2 cells (17). Interestingly, we observed corresponding Nm signals in their *Drosophila* homologs, Top1 and Act5C. However, the precise locations of these Nm sites do not exhibit a strictly conserved positional pattern between the species (**Figure S5A, S5B**). This lack of positional conservation may reflect evolutionary divergence. Alternatively, it could arise from differences in cell culture conditions or developmental stages, mirroring the dynamic regulation of Nm stoichiometry observed in rRNA across varying cellular states (47).

Collectively, these results establish that mRNA 2’-O-methylation is widespread in *Drosophila* S2 cells, could occur through a snoRNA-independent mechanism, and preferentially targets coding regions of functionally related transcripts, revealing a previously uncharacterized regulatory layer of the *Drosophila* mRNA epitranscriptome.

## Discussion

In this study, we combined small RNA sequencing with RibOxi-seq2 analysis to profile snoRNA expression and 2’-O-methylation sites transcriptome-wide in *Drosophila melanogaster* S2 cells. We identified and quantified the expression of 246 snoRNAs that enabled the refinement of existing snoRNA annotations, yielding more precise 5’ and 3’ boundary coordinates. Moreover, RibOxi-seq2 globally mapped Nm modifications across rRNAs, snRNAs, tRNAs, lncRNAs, and mRNAs, substantially expanding the known landscape of Nm-modified transcripts in *Drosophila*. Collectively, these results correlate snoRNA expression with Nm modifications, enabling future studies of snoRNA-guided modification in *Drosophila*.

Our RibOxi-seq2 profiling recovered approximately 94% and 71.4% of previously annotated Nm sites within the 18S and 28S rRNAs in *Drosophila* respectively, consistent with the single-nucleotide resolution reported for this method in prior studies of human and *Trypanosoma brucei (17, 23–25)*. Beyond the annotated sites, we identified and validated one novel Nm modification in the 5.8S rRNA and two novel Nm modifications in the large subunit of mitochondrial rRNA, extending the known repertoire of Nm-modified rRNA positions in *Drosophila*.

Our analysis mapped conserved Nm sites across a broad range of spliceosomal snRNAs, including U2, U4, U4atac, U5, and U6, and we catalogued the corresponding scaRNA guides previously reported to direct each modification (39). Notably, a novel Cm24 site in U5 snRNA was discovered was validated by Nm-VAQ. However, we were unable to identify a scaRNA predicted to target this position, suggesting that this modification may be deposited by an unknown scaRNA. In tRNAs, we identified two recurrent Nm positions conserved across multiple tRNA variants. These positions correspond to well-characterized Nm sites in human and other eukaryotic tRNAs, which are installed by the standalone methyltransferase FTSJ. In addition, RibOxi-seq2 profiling detected m5Um modifications in tRNAs, extending the repertoire of tRNA 2’-O-methylation events captured by RibOxi-seq2. The conservation of both the modification sites and their enzymatic machinery underscores a conserved regulatory machinery for tRNA 2’-O-methylation. Taken together, these findings indicate that both the scaRNA-guided snRNA modification system and the enzyme-directed tRNA modification process are highly conserved between *Drosophila* and humans. This observation is consistent with essential functional roles for Nm modifications in spliceosome assembly, tRNA stability and translational fidelity.

Overall, the near-complete agreement between our profiled Nm sites in rRNAs, tRNAs, and snRNAs and previously published annotations supports the robustness of RibOxi-seq2. Nonetheless, the method has inherent limitations. Most notably, because detection relies on periodate oxidation at 2’-OH groups, closely positioned Nm sites cannot be individually resolved, as the downstream modification blocks access to its upstream neighbor. In addition, other 2’-O-modified bases with similar chemical properties, such as m5Um, can potentially be misidentified as Nm, although such modifications are far less abundant and therefore contribute only a minor fraction of false positives. A promising extension of our approach would be to combine RibOxi-seq2 with bisulfite sequencing, subtracting the Nm signal from bisulfite-derived methylation profiles would allow specific mapping of m5Um and other hybrid modifications, and may represent a useful direction for resolving the distinct contributions of these closely related RNA modifications in future studies.

Using an empirical detection threshold of 500 reads, our RibOxi-seq2 profiling identified Nm sites across 242 lncRNAs in *Drosophila*. The prevalence and functional significance of 2’-O-methylation in lncRNAs remains largely unexplored, and for the sites reported here it is not yet clear whether they are deposited by box C/D snoRNPs, by standalone methyltransferases, or through other enzymatic mechanisms, nor what consequences they have for individual lncRNA function. The systematic characterization provided here therefore serves as a reference annotation of lncRNA Nm sites in *Drosophila* and offers a starting point for future studies addressing the guide dependencies, enzymatic origins, and regulatory roles of 2’-O-methylation in the non-coding RNA biology.

Conservation analysis identified highly conserved Nm sites across rRNA, snRNA, and tRNA species, suggesting that these modifications play evolutionarily conserved roles in RNA folding and function. Interestingly, we also observed cross-species modification divergence, specific Nm sites in *Drosophila* correspond to ac4C modifications in their human homologs. Furthermore, we identified novel modifications within the larger stem-loop regions of snRNAs. Together, these conserved and divergent modification patterns reveal a complex, additional layer of epitranscriptomic regulation.

Interestingly, the data revealed a widespread landscape of 2,057 Nm events in mRNA substrates, a subset of which were independently validated by Nm-VAQ. To our knowledge, this represents the first systematic transcriptome-wide characterization of mRNA Nm modifications in *Drosophila*. The modified positions spanned the 5’ UTR, CDS, and 3’ UTR, with the most significant enrichment within the CDS region. A comparable distribution pattern has previously been reported in several human cell lines, including HEK293, CT2, and H9 neuronal stem cells (16–19), suggesting that the bias toward CDS Nm deposition may be an evolutionarily conserved feature of the mRNA Nm modification. The functional roles of individual mRNA Nm modifications remain to be elucidated, although prior work has implicated these modifications in the regulation of mRNA stability and, more broadly, in fine-tuning translational dynamics. Gene Ontology analysis of Nm-modified mRNAs revealed enrichment for components and cofactors of box C/D snoRNP complexes, which would be consistent with snoRNA-guided deposition. However, computational snoScan searches failed to identify canonical box C/D snoRNA guides predicted to target any of these mRNA sites. This notable paradox raises the possibility that the mRNA Nm modifications in *Drosophila* are installed through non-canonical mechanisms, such as guide-independent methyltransferase activity or snoRNA-guided targeting with base-pairing patterns that diverge from the consensus rules. One intriguing possibility is that many such modifications may be directed by non-canonical interactions involving snoRNA members not previously thought to participate in this activity. Notably, in human cells, evidence has been reported that the abundant box C/D snoRNAs U3 and U8, with known major functions in rRNA maturation, may in fact also direct mRNA modifications. This issue will require experimental identification of the responsible enzymatic machinery and will represent an important direction for future work.

Together, the complementary application of RibOxi-seq2 for transcriptome-wide discovery, Nm-VAQ for site-level validation, and a refined snoRNA atlas for mechanistic inference provides a framework for linking Nm modification sites to their guide RNAs and regulatory contexts in *Drosophila*.

## Material and Methods

### Cell Culture

Schneider’s *Drosophila* Line 2 (S2) cell line was obtained from ATCC (ATCC, CRL-1963) and cultured at 25°C in Schneider’s *Drosophila* Medium (Gibco, R69007) supplemented with 10% FBS (ATCC, 30-2020) and 1% Penicillin-Streptomycin (Gibco, 15140122). Cells were maintained at a density of 0.5 – 2 × 10□ cells/mL and subcultured every 3–4 days.

### RNA Isolation

Total RNA was isolated from cultured S2 cells using the TRIzol. In brief, one million cells were collected and lysed in TRIzol reagent (Invitrogen, #15596026). After incubation at room temperature for 5 minutes, 0.2 mL of chloroform (Invitrogen, #288306) per mL of TRIzol was added, followed by vigorous shaking and centrifugation at 12,000 × g for 15 minutes at 4°C. The aqueous phase containing RNA was collected, mixed with 0.5 mL of isopropanol (Invitrogen, #19030) per mL of TRIzol, incubated for 10 minutes, and centrifuged at 12,000 × g for 10 minutes at 4°C. The resulting RNA pellet was washed with 75% ethanol, air-dried, and resuspended in RNase-free water. The integrity of the isolated mRNA was examined using RNA ScreenTape (Agilent, #5067-5576). mRNA was isolated from 1 – 1.5 mg of total RNA using the NEBNext Poly(A) mRNA Magnetic Isolation Module (New England Biolabs, #E7490) following the manufacturer’s protocol. The integrity of the isolated mRNA was examined using RNA ScreenTape (Agilent, #5067-5576).

### mRNA Truseq library preparation

Illumina TruSeq stranded mRNA libraries (Illumina, 20020594) were prepared in two technical replicates from 500 ng of mRNA isolated from S2 cells, following the manufacturer’s protocol. The mRNA Truseq libraries were pooled and sequenced in paired end manner on the Illumina NextSeq 2000 system.

### Small RNA library preparation

One million S2 cells were lysed with the IB buffer supplied in the Lexogen SPLIT RNA Purification kit (Lexogen, SKU 008.48), followed by phenol-chloroform RNA extraction as per the manufacturer’s instructions. The integrity of the isolated mRNA was examined using RNA ScreenTape (Agilent, #5067-5576). RNA species from S2 cells with size smaller than 200 nt were enriched through spin column purification. Two biological replicates of the small RNA libraries were prepared using the Lexogen Small RNA Library Prep kit (Lexogen, SKU 052.08) following the standard protocol. Additionally, two biological replicates of the small RNA libraries were prepared using the same kit with modifications. After both the A3 and A5 adapter ligation steps, products were cleaned with RNA Clean & Concentrator-5 (Zymo Research, R1013). The ultraMarathonRT enzyme (RNAConnect, R1002L) was replaced for first-strand synthesis. PCR amplification was performed according to the manufacturer’s protocol. The size selection was carried out for both protocols using 1.2× Ampure XP Reagent following the manufacturer’s protocol. The small RNA libraries were pooled equimolarly and sequenced in paired end manner on the Illumina NextSeq 2000 platform.

### RNA-Seq analysis

TruSeq RNA-Seq and small RNA-seq datasets were processed following a standardized pipeline (48). Fastq files were converted using BCL convert v4.2.7. Sequencing adapters were removed and low-quality bases were trimmed using Cutadapt v2.7 (49). Reads shorter than 20 nucleotides or with an average Phred quality score below 20 were discarded. The quality of raw and trimmed reads was assessed with FastQC (v0.12.1). Filtered reads were aligned to the *Drosophila* dm6 reference genome using STAR v2.7.1 (50). Aligned reads were assigned to genes using featureCounts Subread v2.0. (51).

### RibOxi-seq2 library preparation

The RibOxi-seq2 library for *Drosophila* S2 cells was prepared as previously described (17, 23, 24). In brief, mRNA was isolated from 1 – 1.5 mg of total RNA using the NEBNext Poly(A) mRNA Magnetic Isolation Module (New England Biolabs, #E7490).

The integrity of the isolated mRNA was checked using RNA ScreenTape (Agilent, #5067-5576). The RNA was fragmented using benzonase at 95°C for 3 minutes, and incubated on ice for 40 minutes. The fragmented RNA undergoes RNA oxidation, β-Elimination and dephosphorylation reactions to remove excess nucleotides downstream of the 2’-O-methylated nucleotide. The entire oxidation, β-elimination and dephosphorylation process was repeated three additional times. RNA cleanup was performed using the Zymo RNA Clean & Concentrator-25 (Zymo Research, #R1017) after each cycle. The treated RNA was dephosphorylated using T4 PNK and recovered with standard RNA ethanol precipitation. Then the 3’-DNA linker ligation was performed at 16°C for 18 hours using 20 μL of oxidized RNA, 2 μL of a 3’-DNA linker (10 μM) and 4 μL of T4 RNA Ligase 2 truncated KQ (New England Biolabs, #M0373). The free 3’ DNA linkers from the ligation reactions were removed using 15% TBE-Urea gel. The gel was stained with SYBR Gold Nucleic Acid Gel Stain (Invitrogen, #S11494) by incubating it in a 1× TBE staining solution for 15 minutes with gentle shaking. RNA bands were visualized using a Dark Reader Non-UV Transilluminator (Model: DR-46B), and the desired fragment was eluted from the gel with nuclease-free water and recovered by RNA ethanol precipitation as previously described. The 5’-RNA linker ligation was performed using the ligated RNA product, 2 µL of 5’-RNA linker (50 µM) and T4 RNA Ligase 1 (New England Biolabs, #M0204S). Following the 5’-RNA linker ligation, RNA purification was carried out using the Monarch Spin RNA Cleanup kit (New England Biolabs, #T2030) according to the manufacturer’s instructions, and the RNA product was eluted in 20 µL of nuclease-free water. The first strand of cDNA was synthesized at 50°C at 50°C for 1 hour 1 hour with the ligated RNA product, RT primer and ProtoScript II RT polymerase (New England Biolabs, #M0368S). Subsequently, the RNA was degraded with 1 M Sodium Hydroxide (Fisher Chemical, #SS266-1) at 95°C for 15 minutes. The reaction was quenched with 1 M HCl (Fisher Chemical, #A144-500), and the cDNA was purified using 1.8× AMPure XP Reagent (Beckman Coulter, #A63881) following the manufacturer’s protocol. Library amplification was performed with purified cDNA products using D701 and D501 index primers. The PCR conditions for Q5 High-Fidelity DNA Polymerase (New England Biolabs, #M0491) was described in previous protocol, with annealing temperature for the first cycle at 52°C and remaining cycles at 62°C. The amplified library was purified using ethanol precipitation and P-6 columns, followed by size selection using SPRIselect beads (Beckman Coulter, #B23318). RibOxi-seq2 library was sequenced in a paired-end manner on the Illumina NextSeq 2000 platform.

### RibOxi-seq2 oligonucleotides

Oligonucleotide sequences used in RibOxi-seq2 library preparation:

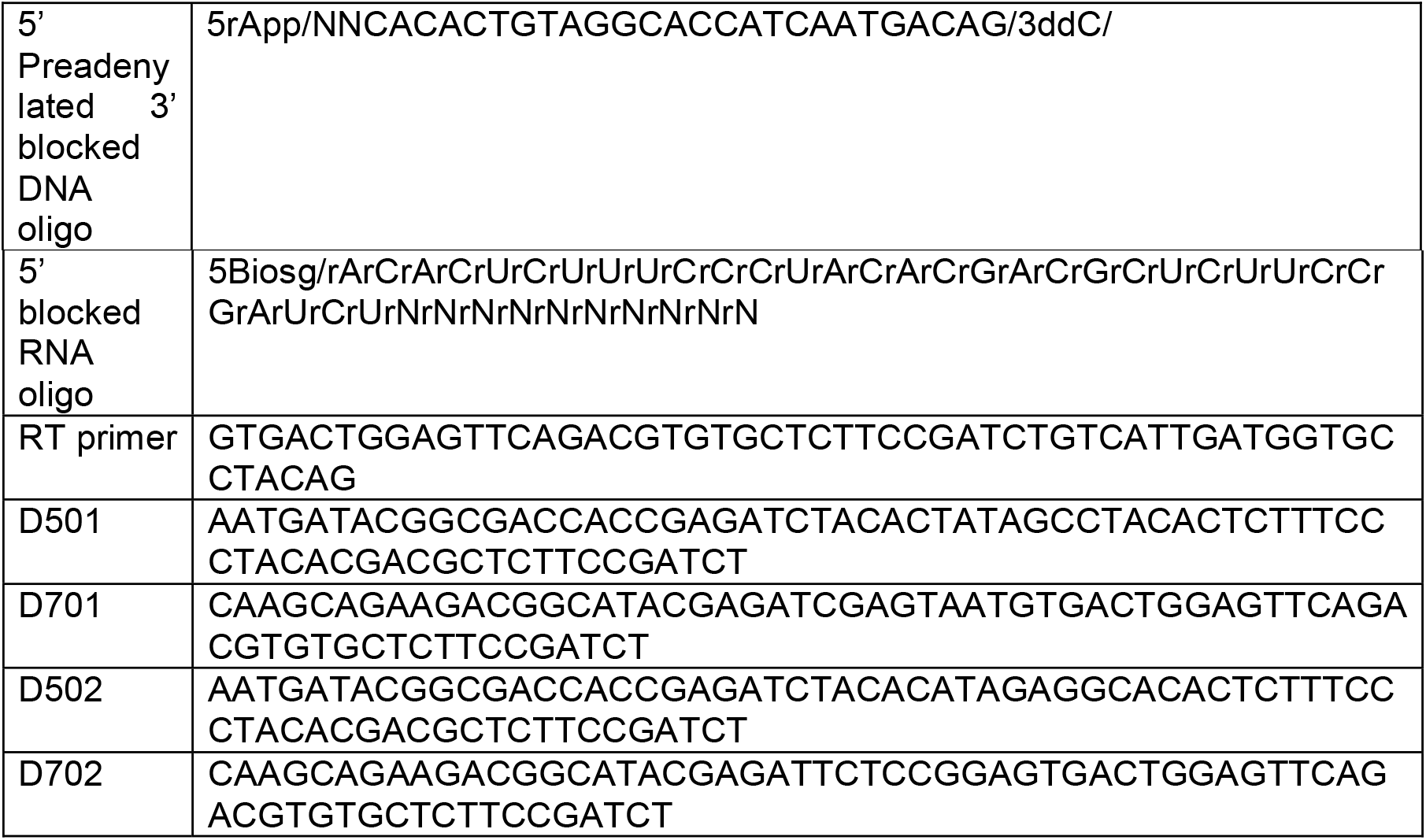

### RibOxi-seq2 analysis

RibOxi-seq2 datasets were analyzed using a modified version of previously reported workflows (https://github.com/yz201906/RibOxi-seq) (17). Sequencing adapters were first trimmed with Cutadapt v2.7, and paired end reads were collapsed into single-end sequences using PEAR v0.9.11 (49, 52). Ten-nucleotide unique molecular identifiers (UMIs) were extracted and appended to the FASTQ headers with the move_umi.py script (17) . Reads shorter than 20 nucleotides or with base quality scores below Q20 were discarded. The remaining high-quality reads were aligned to the reference genome with STAR v2.7.1a (50). Post-alignment processing included filtering and deduplication using UMI-tools v1.1.1 (53). Finally, nucleotide-resolution coverage of 3’ ends, representing putative 2’-O-methylation sites, was computed using Samtools v1.19.2 in combination with Bedtools v2.29.0 (54, 55).

### RNA reference and secondary structure

The *Drosophila melanogaster* 18S rRNA:CR41548, 28S rRNA:CR45844 and snRNA sequences were obtained from RNACentral.org and folded in secondary structure in r2dt (https://rnacentral.org/r2dt) (56, 57).

### Snoscan RNA modification prediction

snoRNA positions were extracted from the dmel-all-r6.54.gtf (https://s3ftp.flybase.org/genomes/Drosophila_melanogaster/dmel_r6.54_FB2023_05/gtf/dmel-all-r6.54.gtf.gz). and dmel-all-r6.58.gtf (https://ftp.flybase.net/genomes/Drosophila_melanogaster/dmel_r6.58_FB2024_03/gtf/dmel-all-r6.58.gtf.gz) (31).

The snoRNA sequences were retrieved from Drosophila melanogaster genome file (https://hgdownload.soe.ucsc.edu/goldenPath/dm6/bigZips/dm6.fa.gz) using Bedtools v2.29.0. Predicted RNA modification sites within these snoRNAs were identified by running snoScan v1.0 against the *Drosophila melanogaster* 18S rRNA:CR41548, 28S rRNA:CR45844, 5.8S rRNA:CR45852 target sequences (57).

### Metagene analysis

Metagene plots summarizing the distribution of Nm sites across mRNA features were generated using metaPlotR (https://github.com/olarerin/metaPlotR) (58).

### Validation and quantification of Nm in RNA

Validation and quantification of 2’-O-methylation sites was performed using the Nm-VAQ method, a previously established RNase H-based assay for both verification and absolute quantification of Nm modifications (19). Chimeric oligonucleotides composed of DNA and 2’-O-methylated RNA, designed to specifically recognize the target Nm site, were synthesized through Integrated DNA Technologies.

For each reaction, 500 ng of total RNA extracted from *Drosophila* S2 cells was combined with 50 pmol of the designed chimera and adjusted to a final volume of 11 µL using 10 µM Tris-HCl buffer (pH 7.0). In cases targeting mRNA specifically, 500 ng of poly(A)-enriched RNA was substituted for total RNA (New England Biolabs, #E7490L). The annealing process involved heating the mixture at 95°C for 1 minute, followed by immediate cooling on ice.

Following annealing, 5 µL of the RNA/chimera solution was incubated with 1 µL of RNase H enzyme, 1 µL of 10× RNase H buffer (New England Biolabs, #B0297S), and 3 µL of nuclease-free water. The remaining 5 µL served as a no-enzyme control and was supplemented with 1 µL of 10× buffer and 4 µL of nuclease-free water. All reactions were incubated at 37°C for 30 minutes.

Subsequent steps diverged depending on RNA abundance. For high-copy RNAs such as rRNA, RNase H-treated samples were subjected to heat inactivation at 90°C for 10 minutes, then chilled on ice. After denaturation, the reaction was diluted 1:5 with nuclease-free water. From this dilution, 1 µL was used for cDNA synthesis using SuperScript III Reverse Transcriptase (Invitrogen, #18080044) and random hexamer (New England Biolabs, #S1230S). qPCR was then conducted with 1 µL of the synthesized cDNA using Power SYBR Green PCR Master Mix (Applied Biosystems, #4368706) on a Bio-Rad CFX96 instrument. Cycling conditions included an initial denaturation at 95°C for 10 minutes, followed by 40 cycles of 95°C for 15 seconds and 60°C for 60 seconds. Specificity of amplification was confirmed through melt curve analysis from 65°C to 95°C in 0.5°C increments, holding 5 seconds at each step.

For low-abundance targets such as mRNA, the 30 µL reaction volume post-RNase H digestion was extracted with 30 µL of phenol:chloroform:isoamyl alcohol (25:24:1) (Millipore Sigma, #77617). The mixture was vortexed vigorously and centrifuged at 12,000 × g for 5 minutes. Approximately 20 µL of the aqueous phase was transferred to a fresh 1.5 mL microcentrifuge tube. RNA purification was carried out using Cytiva Microspin G-50 columns according to the manufacturer’s instructions (Cytiva, #27533001), with elution in 5 µL of nuclease-free water. From this eluate, 1.5 µL was used for cDNA synthesis, and 1 µL of the resulting cDNA was subsequently analyzed by RT-qPCR using the same conditions as described above.

### Nm-VAQ Oligonucleotides

Bold: DNA base; Underlined: 2’-O-methylated RNA base

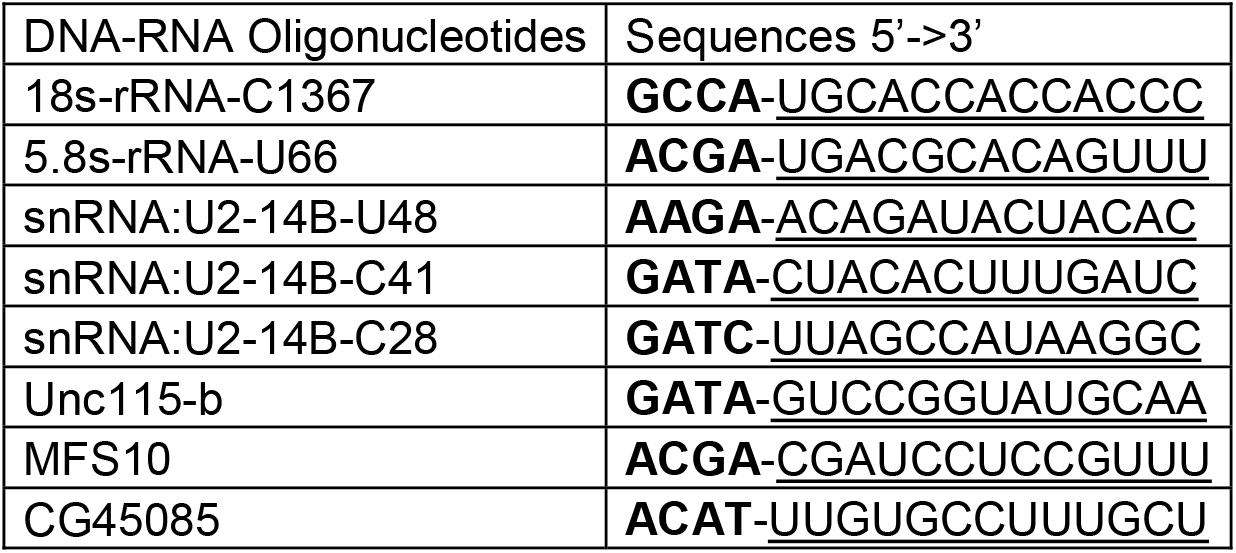

### qPCR Primers

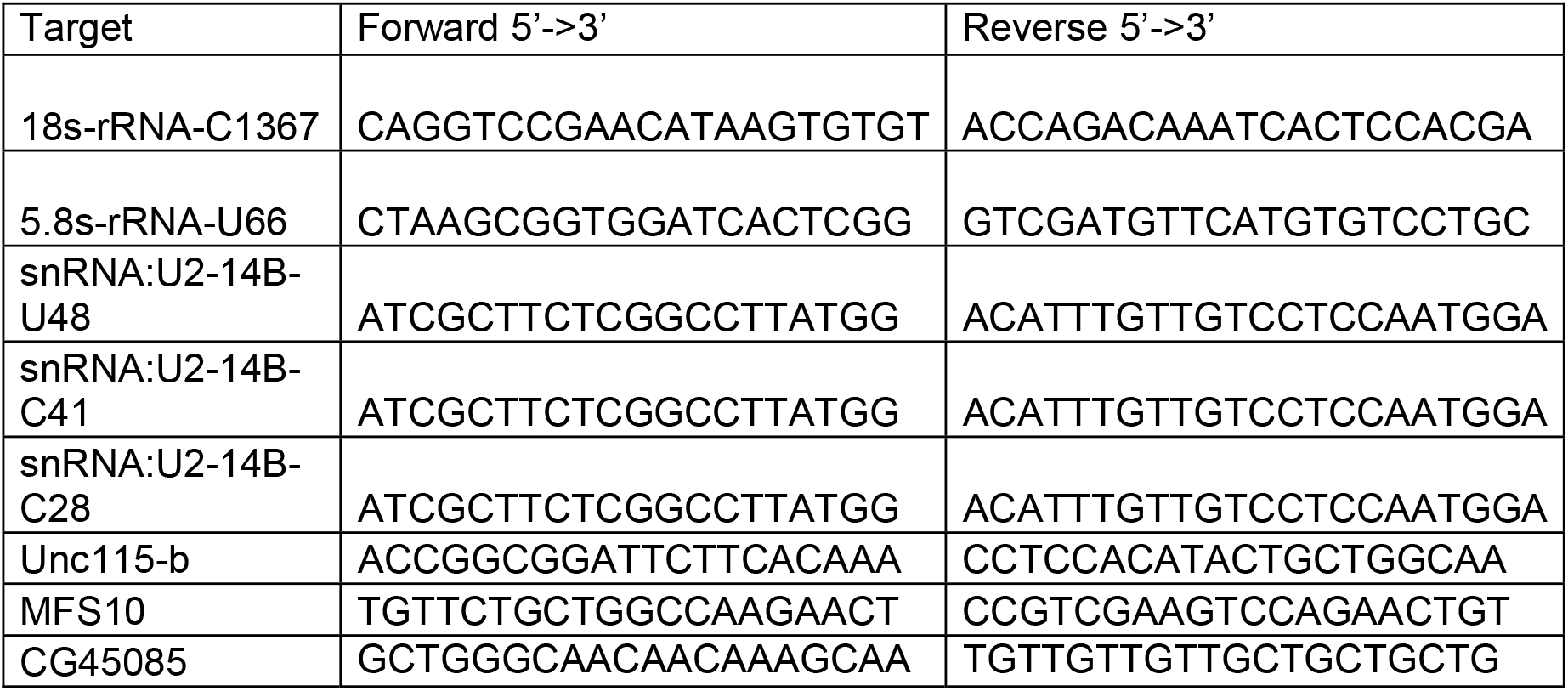

## Data and code availability

Code for analysis is freely available on GitHub at (https://github.com/yz201906/RibOxi-seq) (17). All sequencing data produced in this study have been deposited in the NCBI Gene Expression Omnibus (GEO) under accession number PRJNA1469098.

## Acknowledgments

We thank members of the Graveley and Carmichael labs for discussions. This work was funded by grant R35GM118140 to BRG.

