## Supplementary figures and images for "SnoRNA Expression and RNA 2’-O-Methylation in *Drosophila melanogaster* S2 Cells"

### Supplemental Figure 1

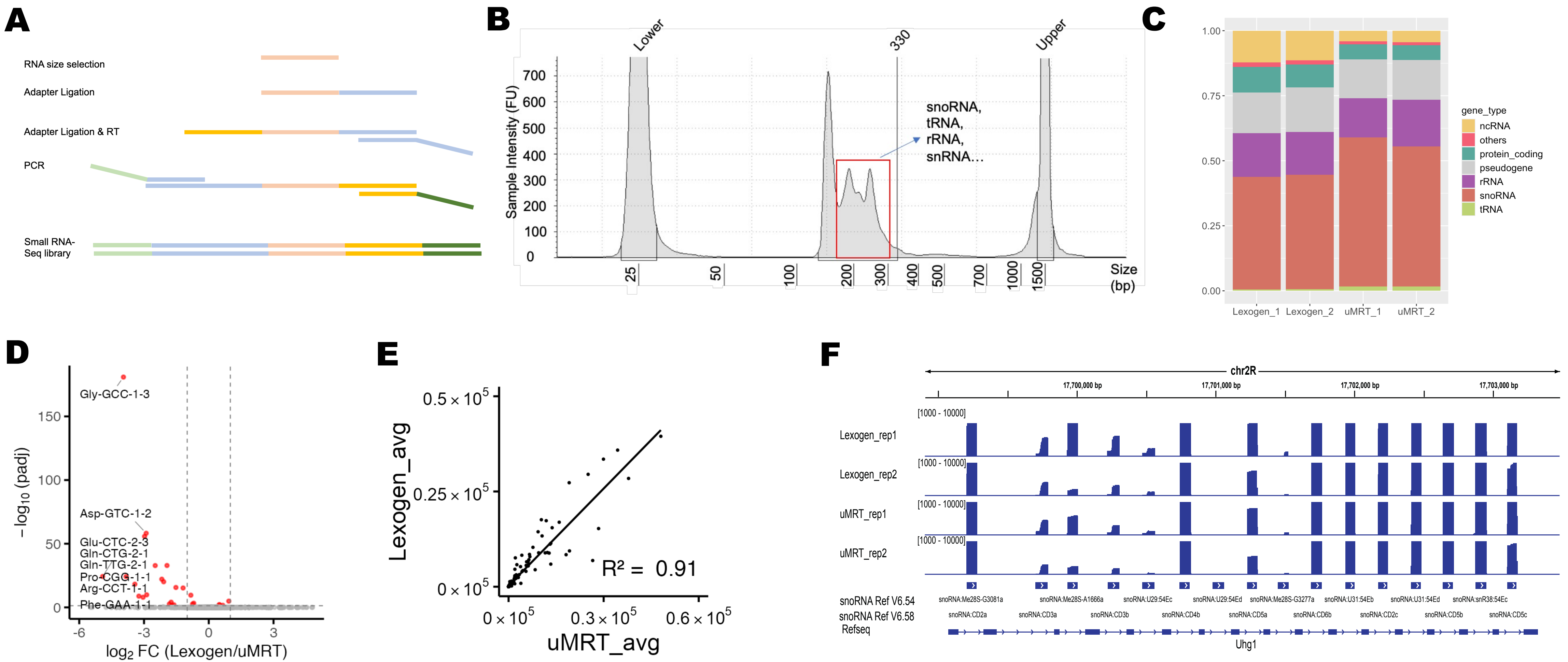

### Supplemental Figure 3

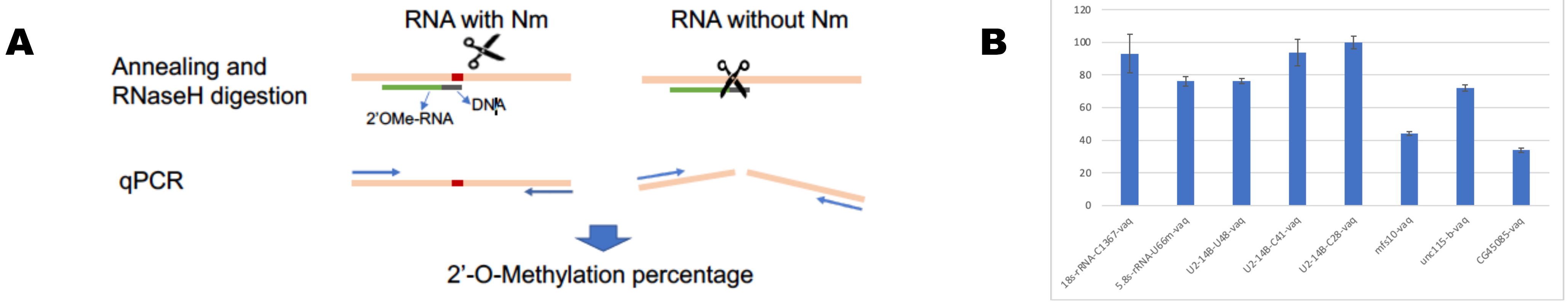

### Supplemental Figure 4

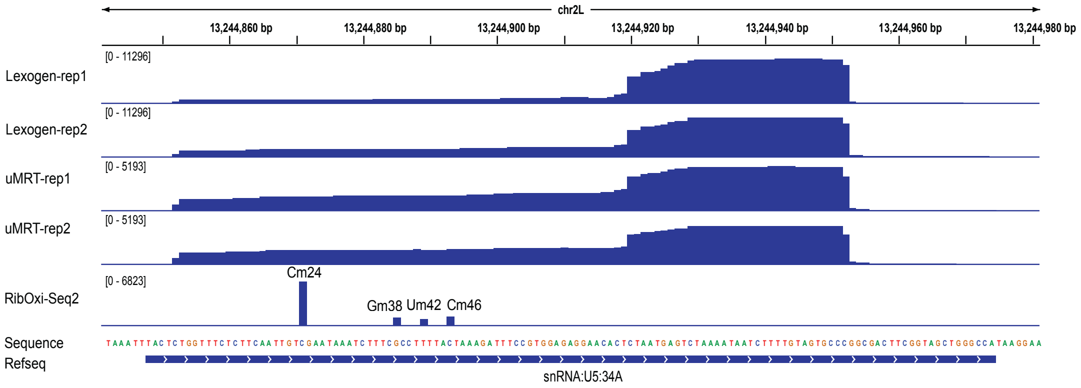

### Supplemental Figure 5

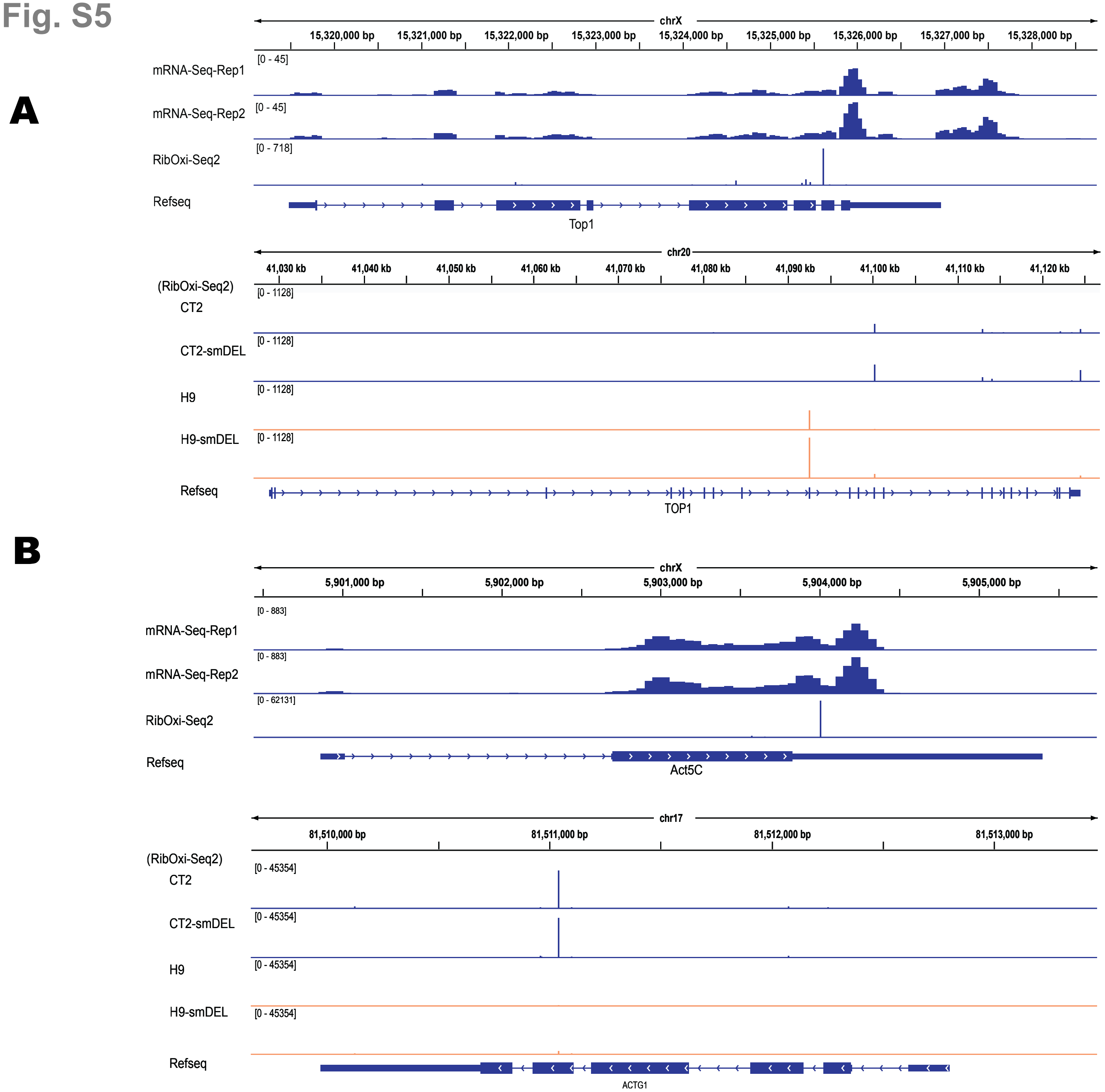
