## Supplemental Figure 2 for "SnoRNA Expression and RNA 2’-O-Methylation in *Drosophila melanogaster* S2 Cells"

**Fig. S2**

**A**

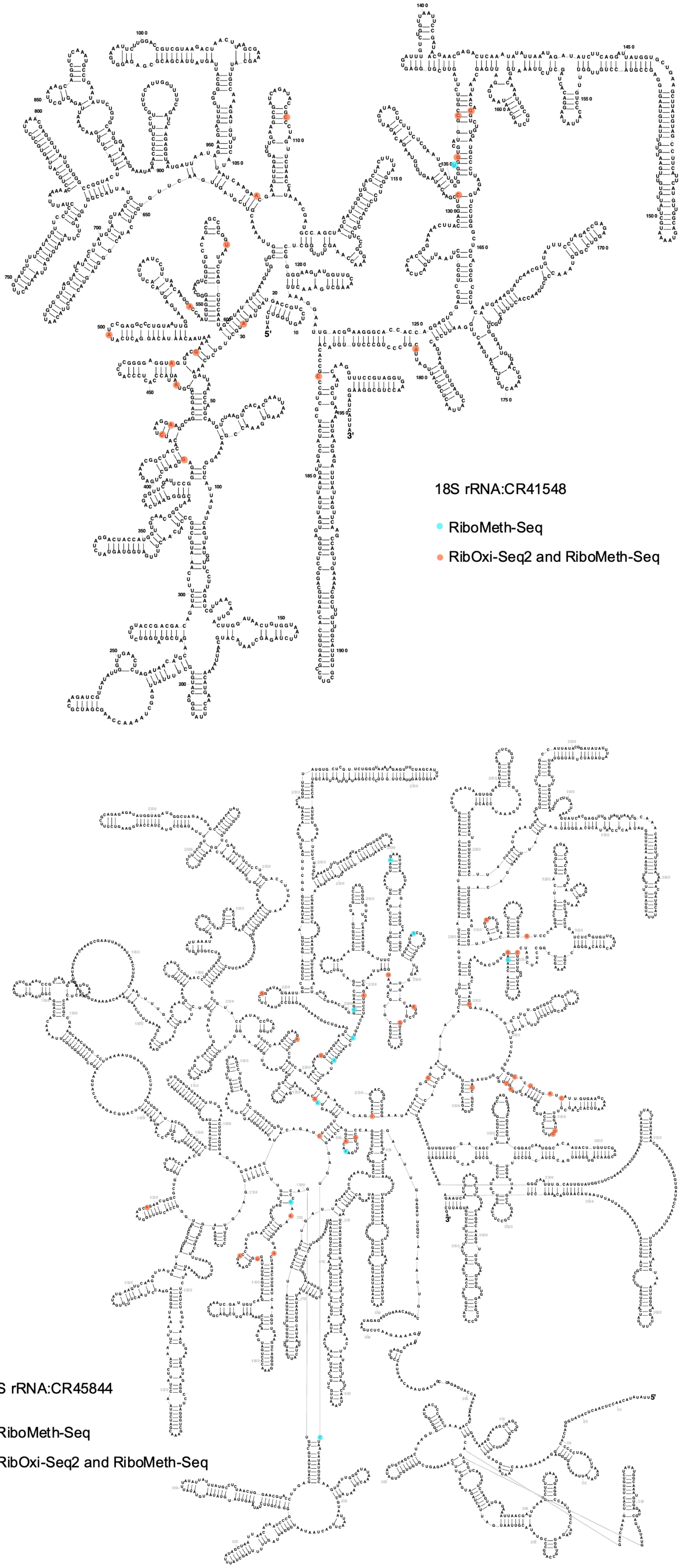
18S rRNA:CR41548
RiboMeth-Seq
RibOxi-Seq2 and RiboMeth-Seq
28S rRNA:CR45844
RiboMeth-Seq
RibOxi-Seq2 and RiboMeth-Seq

#### Fig. S2

**B**

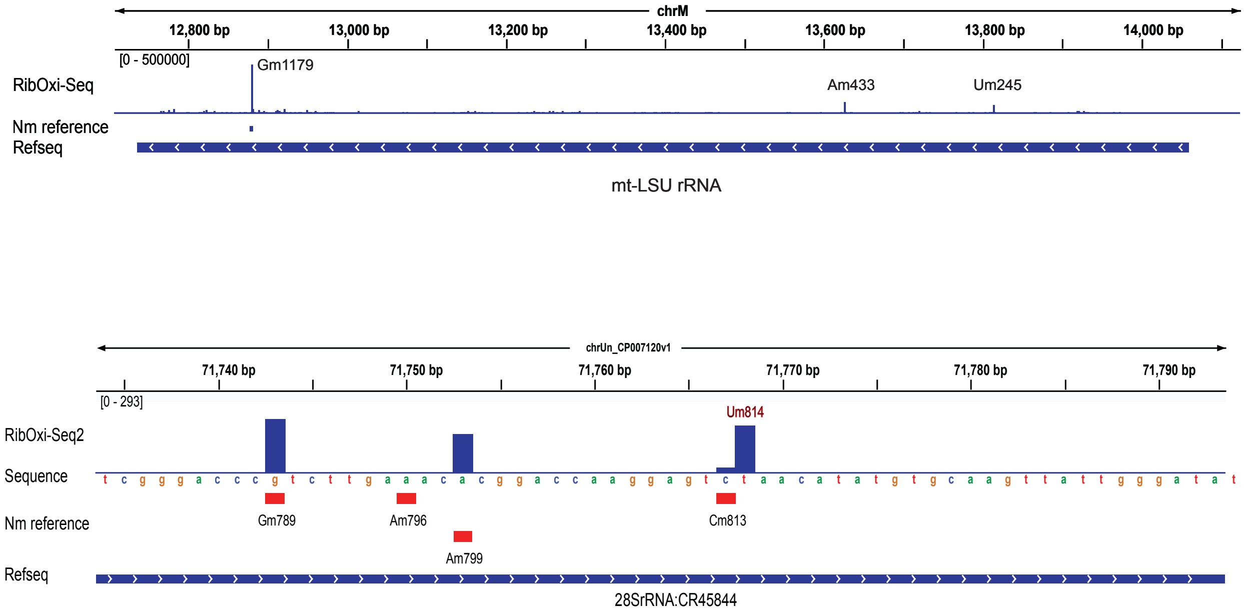

chrM
12,800 bp
13,000 bp
13,200 bp
13,400 bp
13,600 bp
13,800 bp
14,000 bp
[0 - 500000]
RibOxi-Seq
Gm1179
Am433
Um245
Nm reference
Refseq
mt-LSU rRNA
chrUn_CP007120v1
71,740 bp
71,750 bp
71,760 bp
71,770 bp
71,780 bp
71,790 bp
[0 - 293]
RibOxi-Seq2
Um814
Sequence
t c g g g a c c c g t c t t g a a a c a c g g a c c a a g g a g t c t a a c a t a t g t g c a a g t t a t t g g g a t a t
Nm reference
Gm789
Am796
Cm813
Am799
Refseq
28SrRNA:CR45844

### Fig S xx

```
RNA18SN5/1-1869          1 UACCUGGUUGAUCCUGCCAGUAG-CAUAUGCUUGUCUCAAAGAUUAAGCCAUGCAUGUCUGAGUACG 66
18SRNA:CR41548/2-1995    2 UU-CUGGUUGAUCCUGCCAGUAGUUAUAUGCUUGUCUCAAAGAUUAAGCCAUGCAUGUCUAAGUACA 67

RNA18SN5/1-1869         67 CACGGCCGGU-ACAGUGAAACUGCGAAUGGCUCAUUAAAUCAGUUAUGGUUCCUUUGGUCGCU--C- 129
18SRNA:CR41548/2-1995   68 CACGAAU--UAAAAGUGAAACCGCAAAAGGCUCAUUAUAUCAGUUAUGGUUCCUUAGAUCGUUAACA 132

RNA18SN5/1-1869        130 GCUCCUCUCCUACUUGGAUAACUGUGGUAAUUCUAGAGCUAAUACAUGC-----CGACGGGCGCUGA 191
18SRNA:CR41548/2-1995  133 G------U--UACUUGGAUAACUGUGGUAAUUCUAGAGCUAAUACAUGCAAUUAAAACA-----UGA 186

RNA18SN5/1-1869        192 CCCCCUUCGCGGGGGGGAUGCGUGCAUUUAUCA-GAUCAAAACCAACCCGGUCAGCCCCUCUCCGGC 257
18SRNA:CR41548/2-1995  187 ACC--UUAU---GGGACAUG--UGCUUUUAUUAGGCU-AAAACCAA--------------------- 224

RNA18SN5/1-1869        258 CCCGGCCGGGGGGCGGGCGCCGG--CG-GCU-UUGG-UG-ACUCUAGAUAACCUCGGGCCGAUCGCA 318
18SRNA:CR41548/2-1995  225 ------------GCGAUCGCAAGAUCGUUAUAUUGGUUGAACUCUAGAUAACAU---GCAGAUCGUA 276

RNA18SN5/1-1869        319 CGCCCCCCGUGG-C--G---GCGACGAC---CCAUUCGAACGUCUGCCCUAUCAACUUUCGAUGGUA 376
18SRNA:CR41548/2-1995  277 ---------UGGUCUUGUAC-CGACGACAGAUCUUUCAAAUGUCUGCCCUAUCAACUUUUGAUGGUA 333

RNA18SN5/1-1869        377 GUCGCCGU--GCCUACCAUGG-UGACCACGGGUGACGGGGAAUCAGGGUUCGAUUCCGGAGAGGGAG 440
18SRNA:CR41548/2-1995  334 GUAUC--UAGGACUACCAUGGUUG-CAACGGGUAACGGGGAAUCAGGGUUCGAUUCCGGAGAGGGAG 397

RNA18SN5/1-1869        441 CCUGAGAAACGGCUACCACAUCCAAGGAAGGCAGCAGGCGCGCAAAUUACCCACUCCC-GACCCGGG 506
18SRNA:CR41548/2-1995  398 CCUGAGAAACGGCUACCACAUCUAAGGAAGGCAGCAGGCGCGUAAAUUACCCACUCCCAG-CUCGGG 463

RNA18SN5/1-1869        507 GAGGUAGUGACGAAAAAUAACAAUACAGGACUC-U-UUCGAGGCCCUGUAAUUGGAAUGAGUCCACU 571
18SRNA:CR41548/2-1995  464 GAGGUAGUGACGAAAAAUAACAAUACAGGACUCAUAUCCGAGGCCCUGUAAUUGGAAUGAGUACACU 530

RNA18SN5/1-1869        572 UUAAAUCCUUUAACGAGGAUCC-AUUGGAGGGCAAGUCUGGUGCCAGCAGCCGCGGUAAUUCCAGCU 637
18SRNA:CR41548/2-1995  531 UUAAAUCCUUUAACAAGGA-CCAAUUGGAGGGCAAGUCUGGUGCCAGCAGCCGCGGUAAUUCCAGCU 596

RNA18SN5/1-1869        638 CCAAUAGCGUAUAUUAAAGUUGCUGCAGUUAAAAAGCUCGUAGUUGGAUCUUGGG------AGCGGG 698
18SRNA:CR41548/2-1995  597 CCAAUAGCGUAUAUUAAAGUUGUUGCGGUUAAAACGUUCGUAGUUGAA-CUUGUGCUUCAUA-CGGG 661

RNA18SN5/1-1869        699 ------------C----G-GGCG-GUCCG---CC-----G--CG-AGGCG-AGC-CACCGCCCGUCC 734
18SRNA:CR41548/2-1995  662 UAGUACAACUUACAAUUGUGGUUAGUACUAUACCUUUAUGUAUGUAAGCGUA--UUACCG---GUG- 722

RNA18SN5/1-1869        735 CCG----C---------------CCCUUG---CCUCUC----G---GCGCCC---------CCU-CG 762
18SRNA:CR41548/2-1995  723 --GAGUUCUUAUAUGUGAUUAAAUACUUGUAUCUUUUCAUAUGUUC-CUCCUAUUUAAAAACCUGCA 786

RNA18SN5/1-1869        763 --A-UGCUCUU-AGCUGAGUGUCCCGCG-GGGCCCGAAGCGU-UUACUUUGAAAAAAUUAGAGUG-U 822
18SRNA:CR41548/2-1995  787 UUAGUGCUCUUAAAC-GAGUGUUAUU-GUGGGCCGGUA-C-UAUUACUUUGAACAAAUUAGAGUGCU 849

RNA18SN5/1-1869        823 UCAAAGCAGGC-CCGAGCCGCCUGGAUA--C--CGCAGCUAGGAAUAAUGGAAUAGGACCGCGGUUC 884
18SRNA:CR41548/2-1995  850 U-AAAGCAGGCUUCAAAU-GCCUGAAUAUUCUGUGCA--UGGGA-UAAUGAAAUAAGACCUCUGUUC 911

RNA18SN5/1-1869        885 U-AUUUUGUUGGUUUUCGGAAC-UGAGGCCAUGAUUA--AGAGGGACGGCC-GGGGGCAUUCGUAUU 946
18SRNA:CR41548/2-1995  912 UGCUUUCAUUGGUUUUCAGAUCAAGAGGUAAUGAUUAAUAGAAGCA--GUUUGGGGGCAUUAGUAUU 976

RNA18SN5/1-1869        947 GCGCCGCUAGAGGUGAAAUUCUUGGACCGGCGCAAGACGGAC-CAGAGCGAAAGCAUUUGCC-AAGA 1011
18SRNA:CR41548/2-1995  977 ACGACGCGAGAGGUGAAAUUCUUGGACCGUCGUAAGACUAACUUA-AGCGAAAGCAUUUGCCAAAGA 1042

RNA18SN5/1-1869       1012 AUGUUUUCAUUAAUCAAGAACGAAAGUCGGAGGUUCGAAGACGAUCAGAUACCGUCGUAGUUCCGAC 1078
18SRNA:CR41548/2-1995 1043 -UGUUUUCAUUAAUCAAGAACGAAAGUUAGAGGUUCGAAGGCGAUCAGAUACCGCCCUAGUUCUAAC 1108

RNA18SN5/1-1869       1079 CAUAAACGAUGCC-GACCGGCGA-UGCGGCG--GCGUUA-UUCCCAUGACCCGC-CGGGCAGCUU-C 1138
18SRNA:CR41548/2-1995 1109 CAUAAACGAUGCCAG-CUAGCAAUUG-GGUGUAGC--UACUUUU-AUGGCUCUCUCAGUC-GCUUCC 1169

RNA18SN5/1-1869       1139 CGGGAAACCAAAG-UCUUUGGGUUCCGGGGGGAGUAUGGUUGCAAAGCUGAAACUUAAAGGAAUUGA 1204
18SRNA:CR41548/2-1995 1170 CGGGAAACCAAAGCU-UUUGGGCUCCGGGGGAAGUAUGGUUGCAAAGCUGAAACUUAAAGGAAUUGA 1235

RNA18SN5/1-1869       1205 CGGAAGGGCACCACCAGGAGUGGAGCCUGCGGCUUAAUUUGACUCAACACGGGAAACCUCACCCGGC 1271
18SRNA:CR41548/2-1995 1236 CGGAAGGGCACCACCAGGAGUGGAGCCUGCGGCUUAAUUUGACUCAACACGGGAAAACUUACCAGGU 1302

RNA18SN5/1-1869       1272 CCGGACACGGACAG---GAUU-GACAGAUUGAUAGCUCUUUCUCGAUUCCGUGGGUGGUGGUGCAUG 1334
18SRNA:CR41548/2-1995 1303 CCGAACAU--A-AGUGUG-UAAGACAGAUUGAUAGCUCUUUCUCGAAUCUAUGGGUGGUGGUGCAUG 1365

RNA18SN5/1-1869       1335 GCCGUUCUUAGUUGGUGGAGCGAUUUGUCUGGUUAAUUCCGAUAACGAACGAGACUC---------- 1391
18SRNA:CR41548/2-1995 1366 GCCGUUCUUAGUUCGUGGAGUGAUUUGUCUGGUUAAUUCCGAUAACGAACGAGACUCAAAUAUAUUA 1432

RNA18SN5/1-1869       1392 -----------------------------------------------------UGGC------AUGC 1399
18SRNA:CR41548/2-1995 1433 AAUAGAUAUCUUCAGGAUUAUGGUGCUGAAGCUUAUGUAGCCUUCAUUCAUGUUGGCAGUAAAAUGC 1499

RNA18SN5/1-1869       1400 UAACU--------A---GU-----UACG-CGACCCCG-AGCGGUCGG--CGUCCC----------- 1435
18SRNA:CR41548/2-1995 1500 UUAUUGUGUUUGAAUGUGUUUAUGUAAGUGGAGCC--GUACCUGUUGGUUUGUCCCAUUAUAAGGAC 1564

RNA18SN5/1-1869       1436 -CCAACUUCUUAGAGGGACAAGUGGCGU-UCAGC--CACCCGAGAUUGAGCAAUAACAGGUCUGUGA 1498
18SRNA:CR41548/2-1995 1565 ACUAGCUUCUUAAAUGGACAAAUUGCGUCU-AGCAAUAAU-GAGAUUGAGCAAUAACAGGUCUGUGA 1629

RNA18SN5/1-1869       1499 UGCCCUUAGAUGUCCGGGGCUGCACGCGCGCUACACUGACUGGCUCAGCGUGUGCCUA-CCCUACG- 1563
18SRNA:CR41548/2-1995 1630 UGCCCUUAGAUGUCCUGGGCUGCACGCGCGCUACAAUGAAAGUAUCAACGUGUA--UUU-CCUA-GA 1692

RNA18SN5/1-1869       1564 CCG-GCAGG-CGCGGGUAACCCGUUGAACCCCAUUCGUGAUGGGGAUCGGGGAUUGCAAUUAUUCCC 1628
18SRNA:CR41548/2-1995 1693 CCGAG-AGGUC-CGGGUAAACCGCUGAACCACUUUCAUGCUUGGGAUUGUGAACUGAAACUGUUCAC 1757

RNA18SN5/1-1869       1629 CAUGAACGAGGAAUUCCCAGUAAGUGCGGGUCA-UAAGCUUGCGUUGAUUAAGUCCCUGCCCUUUGU 1694
18SRNA:CR41548/2-1995 1758 -AUGAACUUGGAAUUCCCAGUAAGUGUGAGUCAUUAA-CUCGCAUUGAUUACGUCCCUGCCCUUUGU 1822

RNA18SN5/1-1869       1695 ACACACCGCCCGUCGCUACUACCGAUUGGAUGGUUUAGUGAGGCCCUCGGAUCGGCCCCGCCGGGGU 1761
18SRNA:CR41548/2-1995 1823 ACACACCGCCCGUCGCUACUACCGAUUGAAUUAUUUAGUGAGGUC-UC----CGGA-----CGUGAU 1879

RNA18SN5/1-1869       1762 C---G-GCCCACGGCCCU--G-G---CGGA-G---CGCUGAGAA---GACGGUCGAACUUGACUAUC 1811
18SRNA:CR41548/2-1995 1880 CACUGUG---ACG-CCUUGCGUGUUACGGUUGUUUCGC--AAAAGUUGAC---CGAACUUGAUUAUU 1937
```

| | | | |
|---|---|---|---|
| *RNA18SN5/1-1869* | 1812 | UAGAGGAAGUAAAAGUCGUAACAAGGUUUCCGUAGGUGAACCUGCGGAAGGAUCAUUA | 1869 |
| *18SRNA:CR41548/2-1995* | 1938 | UAGAGGAAGUAAAAGUCGUAACAAGGUUUCCGUAGGUGAACCUGCGGAAGGAUCAUUA | 1995 |

Fig S xxx

```
RNA28SN5/1-5070           1 -----CGCGACCUCAGAUCAGACGUGGCGACCCGCUGAAUUUAAGCAUAUUAGUCAGCGGAGGAGA 61
28SRNA:CR45844/1-3970     1 UUAUAUACAACCUCAACUCAUAUGGGACUACCCCCUGAAUUUAAGCAUAUUAAUUAGGGGAGGAAA 66

RNA28SN5/1-5070          62 AGAAACUAACCAGGAUUCCCUCAGUAACGGCGAGUGAACAGGGAAGAGCCCAGCGCCGAAUCCCCG 127
28SRNA:CR45844/1-3970    67 AGAAACUAACAAGGAUUUUCUUAGUAGCGGCGAGCGAAAAGAAAACAGUUCAGCACUAAGUCACUU 132

RNA28SN5/1-5070         128 CCCCGCGGCGGGGCGCGGGACAUGUGGCGUACGGAAGACCCGCUCCCCGGCGCCGCUCGUGGGGGG 193
28SRNA:CR45844/1-3970   133 UGUCUAUAUGGCAAAUGUGAGAUGCAGUGUAUGGAGCGUCAAUAUUCUAGUAUGAGAAAUUAACGA 198

RNA28SN5/1-5070         194 CCCAAGUCCUUCUGAUC-GAGGCCC--AGCCCGUGGACGGUGUGAGGCCGGUAGCGGCCCCCGGCG 256
28SRNA:CR45844/1-3970   199 UUUAAGUCCUUCUUAAAUGAGGCCAUUUACCCAUAGAGGGUGCCAGGCCCGUAUAACGUUAAUGAU 264

RNA28SN5/1-5070         257 CGCCGGGCCCGGGUCUUCCCGGAGUCGGGUUGCUUGGGAAUGCAGCCCAAAGCGGGUGGUAAACUC 322
28SRNA:CR45844/1-3970   265 U-ACUAGAUGAUGUUUCCAAAGAGUCGUGUUGCUUGAUAGUGCAGCACUAAGUGGGUGGUAAACUC 329

RNA28SN5/1-5070         323 CAUCUAAGGCUAAAUACCGGCACGAGACCGAUAGUCAACAAGUACCGUAAGGGAAAGUUGAAAAGA 388
28SRNA:CR45844/1-3970   330 CAUCUAAAACUAAAUAUAACCAUGAGACCGAUAGUAAACAAGUACCGUGAGGGAAAGUUGAAAAGA 395

RNA28SN5/1-5070         389 ACUUUGAAGAGAGAGUUCAAGAGGGCGUGAAACCGUUAAGAGGUAAACGGGUGGGGUCCGCGCAGU 454
28SRNA:CR45844/1-3970   396 ACUCUGAAUAGAGAGUUAAACAGUACGUGAAACUGCUUAGAGGUUAAGCCCGAUGAACCUGAAUAU 461

RNA28SN5/1-5070         455 CCG-CCCGGAGGAUUCAACCCGGCGGCGGGUCCGGCCGUGUCGGCGGCCCGGCGGAUCUUUCCCGC 519
28SRNA:CR45844/1-3970   462 CCGUUAUGGAAAAUUCAUCAUUAAAA-----------UUG-------------UAAUAUUUAA--- 500

RNA28SN5/1-5070         520 CCCCCGUUCCUCCCGACCCCUCCACCCGCCCUCCCUUCCCCCGCCGCCCCUCCUCCUCCUCCCCGG 585
28SRNA:CR45844/1-3970   501 ----------------------------------------------------AUAAUAUUAUGAGA 514

RNA28SN5/1-5070         586 AGGGGGCGGGCUCCGGCGGGUGCGGGGGUGGGCGGGCGGGGCCGGGGGUGGGGUCGGCGGGGGACC 651
28SRNA:CR45844/1-3970   515 A----------------------------------------------------------------- 515

RNA28SN5/1-5070         652 GUCCCCCGACCGGCGACCGGCCGCCGCCGGGCGCAUUUCCACCGCGGCGGUGCGCCGCGACC---- 713
28SRNA:CR45844/1-3970   516 --------------------------UAGUGUGCAUUUUUUCCAUAUAAGGACAUUGUAAUCUAUU 555

RNA28SN5/1-5070         714 ---GGCUCCGGGACGGCUGGGAAGGCCCGGCGGGGAAGGUGGCUCGGGGGGCCCCGUCCGUCCGUC 776
28SRNA:CR45844/1-3970   556 AGCAUAUACCAAAUUUAUC------------AUAAAAUAUAACUUAUA------------------ 591

RNA28SN5/1-5070         777 CGUCCGUCCUCCUCCUCCCCCGUCUCCGCCCCCCGGCCCCGCGUCCUCCCUCGGGAGGGCGCGCGG 842
28SRNA:CR45844/1-3970   592 ----------------------GUUU------------------AUUCCAAUUAAAUUGCUUG--- 614

RNA28SN5/1-5070         843 GUCGGGGCGGCGGCGGCGGCGGCGGUGGCGGCGGCGGCGGCGGCGGGACCGAAACCCCCCCCG 908
28SRNA:CR45844/1-3970   615 -------------------------------------------------CAUUUUAACACAGAAUA 631

RNA28SN5/1-5070         909 AGUGUUACAGCCCCCCCGGCAGCAGCACUCGCCGAAUCCCGGGGCCGAGGGAGCGAGACCCGUCGC 974
28SRNA:CR45844/1-3970   632 AAUGUUAUUA----------------AUUUGAUAAA-----GUGCUGAUAGAUUUAUAUGA----- 671

RNA28SN5/1-5070         975 CGCGCUCUCCCCCCUCCCGGCGCCCACCCCCGCGGGGAAUCCCCCGCGAGGGGGGUCUCCCCCGCG 1040
28SRNA:CR45844/1-3970   672 ---------------UUACAGUG----------CG-UUAAUUUUC--------------------- 691

RNA28SN5/1-5070        1041 GGGGCGCGCCGGCGUCUCCUCGUGGGGGGGCCGGGCCACCCCUCCCACGGCGCGACCGCUCUCCCA 1106
28SRNA:CR45844/1-3970   692 -----------GGAAU------------------------UAUAUAAUGGCAU------------- 709

RNA28SN5/1-5070        1107 CCCCUCCUCCCCGCGCCCCCGCCCCGGCGACGGGGGGGGUGCCGCGCGCGGGUCGGGGGGCGGGGC 1172
28SRNA:CR45844/1-3970       ------------------------------------------------------------------

RNA28SN5/1-5070        1173 GGACUGUCCCCAGUGCGCCCCGGGCGGGUCGCGCCGUCGGGCCCGGGGGAGGUUCUCUCGGGGCCA 1238
28SRNA:CR45844/1-3970   710 -AAUUAU---CAUUGAU---------------------------------UUUUGUGUUUAUUAUA 738

RNA28SN5/1-5070        1239 CGCGCGCGUCCCCCGAAGAGGGGGACGGCGGAGCGAGCGCACGGGGUCGGCGGCGACGUCGGCUAC 1304
28SRNA:CR45844/1-3970   739 UGCACUUGUAUGAUUAACAAUGCGA---------------------------AAGAUUCAGGAUAC 777

RNA28SN5/1-5070        1305 CCACCCGACCCGUCUUGAAACACGGACCAAGGAGUCUAACACGUGCGCGAGUCGGGGGCUCGCACG 1370
28SRNA:CR45844/1-3970   778 CUUCGGGACCCGUCUUGAAACACGGACCAAGGAGUCUAACAUAUGUGCAAGUUAUUGGG----AU- 838

RNA28SN5/1-5070        1371 AAAGCCGCCGUGGCGCAAUGAAGGUGAAGGCCGGCGCGCUCGCCGGCCGAGG-------------- 1422
28SRNA:CR45844/1-3970   839 AUAAACCUAAUAGCGUAAUUAACUUGACUAAUAAUGGGAUUAGUUUUUUAGCUAUUUAUAGCUAAU 904

RNA28SN5/1-5070        1423 ---UGGGAUCCCGAGGCCUC---UCCAGUCCG-CCGAGGGCGCACCACCGGCCCGUCUCGCCCGCC 1481
28SRNA:CR45844/1-3970   905 UAACACAAUCCCGGGGCGUUCUAUAUAGUUAUGUAUAAUGUAUAUU--UAUAUUAUUUA---UGCC 965

RNA28SN5/1-5070        1482 GC-GCCGGGGAGGUGGAGCACGAGCGCACGUGUUAGGACCCGAAAGAUGGUGAACUAUGCCUGGGC 1546
28SRNA:CR45844/1-3970   966 UCUAACUGGAA--CGUACCUUGAGCAUAUAUGCUGUGACCCGAAAGAUGGUGAACUAUACUUGAUC 1029

RNA28SN5/1-5070        1547 AGGGCGAAGCCAGAGGAAACUCUGGUGGAGGUCCGUAGCGGUCCUGACGUGCAAAUCGGUCGUCCG 1612
28SRNA:CR45844/1-3970  1030 AGGUUGAAGUCAGGGGAAACCCUGAUGGAAGACCGAAACAGUUCUGACGUGCAAAUCGAUUGUCAG 1095

RNA28SN5/1-5070        1613 ACCUGGGUAUAGGGGCGAAAGACUAAUCGAACCAUCUAGUAGCUGGUUCCCUCCGAAGUUUCCCUC 1678
28SRNA:CR45844/1-3970  1096 AAUUGAGUAUAGGGGCGAAAGACCAAUCGAACCAUCUAGUAGCUGGUUCCUUCCGAAGUUUCCCUC 1161

RNA28SN5/1-5070        1679 AGGAUAGCUGGCGCUCUCGCAGACCCGACGCACCCCCGCCACGCAGUUUUAUCCGGUAAAGCGAAU 1744
28SRNA:CR45844/1-3970  1162 AGGAUAGCUGGUGCAUUUUAAUAUUAUAU----------AAAAUAAUCUUAUCUGGUAAAGCGAAU 1217

RNA28SN5/1-5070        1745 GAUUAGAGGUCUUGGGGCCGAAACGAUCUCAACCUAUUCUCAAACUUUAAAUGGGUAAGAAGCCCG 1810
28SRNA:CR45844/1-3970  1218 GAUUAGAGGCCUUAGGGUCGAAACGAUCUUAACCUAUUCUCAAACUUUAAAUGGGUAAGAACCUUA 1283

RNA28SN5/1-5070        1811 GCUCGCUGGCG-UGGAGCCG--GGCGUGGAAUGCGAGUGCCUAGUGGGCCACUUUUGGUAAGCAGA 1873
28SRNA:CR45844/1-3970  1284 ACUUUCUUGAUAUGAAGUUCAAGGUUAUGAUAUAAUGUGCCCAGUGGGCCACUUUUGGUAAGCAGA 1349

RNA28SN5/1-5070        1874 ACUGGCGCUGCGGGAUGAACCGAACGCCGGGUUAAGGCGCCCGAUGCCGACGCUCAU-CAGACCCC 1938
28SRNA:CR45844/1-3970  1350 ACUGGCGCUGUGGGAUGAACCAAACGUAAUGUUACGGUGCCCAAAUUAACAACUCAUGCAGAUACC 1415

RNA28SN5/1-5070        1939 AGAAAAGGUGUUGGUUGAUAUAGACAGCAGGACGGUGGCCAUGGAAGUCGGAAUCCGCUAAGGAGU 2004
28SRNA:CR45844/1-3970  1416 AUGAAAGGCGUUGGUUGCUUAAAACAGCAGGACGGUGAUCAUGGAAGUCGAAAUCCGCUAAGGAGU 1481
```

| | | | |
|---|---|---|---|
| *RNA28SN5/1-5070* | 2005 | GUGUAACAACUCACCUGCCGAAUCAACUAGCCCUGAAAAUGGAUGGCGCUGGAGCGUCGGCCCAU | 2070 |
| *28SRNA:CR45844/1-3970* | 1482 | GUGUAACAACUCACCUGCCGAAGCAACUAGCCCUUAAAAUGGAUGGCGCUUAAGUUGUAUACCUAU | 1547 |
| *RNA28SN5/1-5070* | 2071 | ACCCGGCCGUCGCCGGCAGUCGAGAGUGGACGGGAGCGGCGGGGGCGGCGCGCGCGCGCGCGCGUG | 2136 |
| *28SRNA:CR45844/1-3970* | 1548 | ACAUUAC------CGCU-----AAAGUAGAUGAUUUA--------------------UAUUACUUG | 1582 |
| *RNA28SN5/1-5070* | 2137 | UGGUGUGCGUCGGAGGGCGGCGGCGGCGGCGGCGGCGGGGGUGUGGGGUCCUUCCCCCGCCCCCCC | 2202 |
| *28SRNA:CR45844/1-3970* | 1583 | UGAUAUAAAUUU------------------------------------------------------ | 1594 |
| *RNA28SN5/1-5070* | 2203 | CCCCACGCCUCCUCCCCUCCUCCCGCCCACGCCCCGCUCCCCGCCCCCGGAGCCCCGCGGACGCUA | 2268 |
| *28SRNA:CR45844/1-3970* | 1595 | ----------------------------------------------------------UGA----- | 1597 |
| *RNA28SN5/1-5070* | 2269 | CGCCGCGACGAGUAGGAGGGCCGCUGCGGUGAGCCUUGAAGCCUAGGGCGCGGGCCCGGGUGGAGC | 2334 |
| *28SRNA:CR45844/1-3970* | 1598 | AACUUUAGUGAGUAGGAAGGU-ACAAUGGUAUGCGUAGAAGUGUUUGGCGUAAGCCUGCAUGGAGC | 1662 |
| *RNA28SN5/1-5070* | 2335 | CGCCGCAGGUGCAGAUCUUGGUGGUAGUAGCAAAUAUUCAAACGAGAACUUUGAAGGCCGAAGUGG | 2400 |
| *28SRNA:CR45844/1-3970* | 1663 | UGCCAUUGGUACAGAUCUUGGUGGUAGUAGCAAAUAAUCGAAUGAGACCUUGGAGGACUGAAGUGG | 1728 |
| *RNA28SN5/1-5070* | 2401 | AGAAGGGUUCCAUGUGAACAGCAGUUGAACAUGGGUCAGUCGGUCCUGAGAGAUGGGCGAGCGCCG | 2466 |
| *28SRNA:CR45844/1-3970* | 1729 | AGAAGGGUUUCGUGUGAACAGUGGUUGAUCACGAGUUAGUCGGUCCUAAGUUCAAGGCGAAAGCCG | 1794 |
| *RNA28SN5/1-5070* | 2467 | U----UCCGAAGGGACGGGCGAUGGCCU----------------------------CCGUUGCCCU | 2500 |
| *28SRNA:CR45844/1-3970* | 1795 | AAAAUUUUCAAGUAAAACAAAAAUGCCUAACUAUAUAAACAAAGCGAAUUAUAAUACACUUGAAUA | 1860 |
| *RNA28SN5/1-5070* | 2501 | CGGCCGAUCGAAAGGGAGUCGGGUUCAGAUCCCCGAAUCCGG--AGUGGC-----------GGAGA | 2553 |
| *28SRNA:CR45844/1-3970* | 1861 | AUUUUGAACGAAAGGGAAUACGGUUCCAAUUCCGUAACCUGUUGAGUAUCCGUUUGUUAUUAAAUA | 1926 |
| *RNA28SN5/1-5070* | 2554 | UGGGCGCCGCGAGGCGUCCAGUGCGGUAACGCGACCGAUCCCGGAGAAGCCGGCGGGAGCCCCGGG | 2619 |
| *28SRNA:CR45844/1-3970* | 1927 | UGGGCCUCG-----UGCUCAUCCUGGCAACAGGAACGACCAUAAAGAAGCCGUCGAGAGAUAUCGG | 1987 |
| *RNA28SN5/1-5070* | 2620 | GAGAGUUCUCUUUUCUUUGUGAAGGGCAGGGCGCCCUGGAAUGGGUUCGCCCCGAGAGAGGGGCCC | 2685 |
| *28SRNA:CR45844/1-3970* | 1988 | AAGAGUUUUCUUUUCUGUUUUAUAGCCGUACUACCAUGGAAGUCUUUCGCAGAGAGAUAUGGUAGA | 2053 |
| *RNA28SN5/1-5070* | 2686 | GUGCCUUGGAAAGCGUCGCGGUUCCGGCGGCGUCCGGUGAGCUCUCGCUGGCCCUUGAAAAUCCGG | 2751 |
| *28SRNA:CR45844/1-3970* | 2054 | UGGGCUAGAAGAGCAUGACAUAUACUGUUGUGUC-GAUAUUUUCUCCUCGGACCUUGAAAAUUUAU | 2118 |
| *RNA28SN5/1-5070* | 2752 | GGG-AGAGGGUGUAAAUCUCGCGCCGGGCCGUACCCAUAUCCGCAGCAGGUCUCCAAGGUGAACAG | 2816 |
| *28SRNA:CR45844/1-3970* | 2119 | GGUGGGGACACGCAAACUUCUCAACAGGCCGUACCAAUAUCCGCAGCUGGUCUCCAAGGUGAAGAG | 2184 |
| *RNA28SN5/1-5070* | 2817 | CCUCUGGCAUGUUGGAACAAUGUAGGUAAGGGAAGUCGGCAAGCCGGAUCCGUAACUUCGGGAUAA | 2882 |
| *28SRNA:CR45844/1-3970* | 2185 | UCUCUAGUC-GAUAGAAUAAUGUAGGUAAGGGAAGUCGGCAAAUUAGAUCCGUAACUUCGGGAUAA | 2249 |
| *RNA28SN5/1-5070* | 2883 | GGAUUGGCUCUAAGGGCUGGGUCGGUCGGGCUGGGGCGCGAAGCGGGGCUGGGCGCGCGCCGCGGC | 2948 |
| *28SRNA:CR45844/1-3970* | 2250 | GGAUUGGCUCUGAAGAUUGAGAUAGUCGGGCUUGAUUGGGAAACAAUAACA--------------- | 2300 |
| *RNA28SN5/1-5070* | 2949 | UGGACGAGGCGCCGCCGCCCCCCCACGCCCGGGGCACCCCCCUCGCGGCCCUCCCCCGCCCCACC | 3014 |
| *28SRNA:CR45844/1-3970* | 2301 | UGGUUUAUGUGC--------------UCGUUCUGGGUAAA----UAG------------------- | 2329 |
| *RNA28SN5/1-5070* | 3015 | CCGCGCGCGCCGCUCGCUCCCUCCCCGCCCCGCGCCCUCUCUCUCUCUCUCUCCCCCGCUCCCCGU | 3080 |
| *28SRNA:CR45844/1-3970* | 2330 | -------AGUUUCUA------------------G-CAUUUAU----GUUAGUUACUUGUU------ | 2359 |
| *RNA28SN5/1-5070* | 3081 | CCUCCCCCCUCCCCGGGGGAGCGCCGCGUGGGGGCGGCGGCGGGGGAGAAGGGUCGGGGCGGCAG | 3146 |
| *28SRNA:CR45844/1-3970* | 2360 | -----------CCCCG-GAUAGUUU--A-------------------------------------- | 2373 |
| *RNA28SN5/1-5070* | 3147 | GGGCCGGCGGCGGCCCGCCGCGGGGCCCCGGCGGCGGGGGCACGGUCCCCCGCGAGGGGGGCCCGG | 3212 |
| *28SRNA:CR45844/1-3970* |  | ------------------------------------------------------------------ |  |
| *RNA28SN5/1-5070* | 3213 | GCACCCGGGGGGCCGGCGGCGGCGGCGACUCUGGACGCGAGCCGGGCCCUUCCCGUGGAUCGCCCC | 3278 |
| *28SRNA:CR45844/1-3970* | 2374 | ---------------GUUACGUAGCCAAUUGUGGAA-----------CUUUCUUG----------- | 2402 |
| *RNA28SN5/1-5070* | 3279 | AGCUGCGGCGGGCGUCGCGGCCGCCCCCGGGGAGCCCGGCGGGCGCCGGCGCGCCCCCCCCCCCAC | 3344 |
| *28SRNA:CR45844/1-3970* | 2403 | --CU-------------------------------------------------------------- | 2404 |
| *RNA28SN5/1-5070* | 3345 | CCCACGUCUCGUCGCGCGCGCGUCCGCUGGGGGCGGGGAGCGGUCGGGCGGCGGCGGUCGGCGGGC | 3410 |
| *28SRNA:CR45844/1-3970* | 2405 | --AAAAUUUUUAA------GAAUACUAU---------------UUGGG------------------ | 2429 |
| *RNA28SN5/1-5070* | 3411 | GGCGGGGCGGGGCGGUUCGUCCCCCCGCCCUACCCCCCCGGCCCCGUCCGCCCCCCGUUCCCCCCU | 3476 |
| *28SRNA:CR45844/1-3970* | 2430 | ------UUAAACCAAUUAGU--------------------------------------U----CUU | 2447 |
| *RNA28SN5/1-5070* | 3477 | CCUCCUCGGCGCGCGGCGGCGGCGGCGGCAGGCGGCGGAGGGGCCGCGGGCCGGUCCCCCCCGCCG | 3542 |
| *28SRNA:CR45844/1-3970* | 2448 | A----------------------------------------------------------------- | 2448 |
| *RNA28SN5/1-5070* | 3543 | GGUCCGCCCCCGGGGCCGCGGUUCCGCGCGGCGCCUCGCCUCGGCCGGCGCCUAGCAGCCGACUUA | 3608 |
| *28SRNA:CR45844/1-3970* | 2449 | ---------------------UUA----------AUUAUAACGAUUAUCAAUUAACAAUCAAUUCA | 2483 |
| *RNA28SN5/1-5070* | 3609 | GAACUGGUGCGGACCAGGGGAAUCCGACUGUUUAAUUAAAACAAAGCAUCGCGAAGGCCCGCGGCG | 3674 |
| *28SRNA:CR45844/1-3970* | 2484 | GAACUGGCACGGACUUGGGGAAUCCGACUGUCUAAUUAAAACAAAGCAUUGUGAUGGCCCU-AGCG | 2548 |
| *RNA28SN5/1-5070* | 3675 | GGUGUUGACGCGAUGUGAUUUCUGCCCAGUGCUCUGAAUGUCAAAGUGAAGAAAUUCAAUGAAGCG | 3740 |
| *28SRNA:CR45844/1-3970* | 2549 | GGUGUUGACACAAUGUGAUUUCUGCCCAGUGCUCUGAAUGUCAAAGUGAAGAAAUUCAAGUAAGCG | 2614 |
| *RNA28SN5/1-5070* | 3741 | CGGGUAAACGGCGGGAGUAACUAUGACUCUCUUAAGGUAGCCAAAUGCCUCGUCAUCUAAUUAGUG | 3806 |
| *28SRNA:CR45844/1-3970* | 2615 | CGGGUCAACGGCGGGAGUAACUAUGACUCUCUUAAGGUAGCCAAAUGCCUCGUCAUCUAAUUAGUG | 2680 |
| *RNA28SN5/1-5070* | 3807 | ACGCGCAUGAAUGGAUGAACGAGAUUCCCAC-UGUCCCUACCUACUAUCCAGCGAAACCACAGCCA | 3871 |
| *28SRNA:CR45844/1-3970* | 2681 | ACGCGCAUGAAUGGAUUAACGAGAUUCCUACUUGUCCCUAUCUACUAUCUAGCGAAACCACAGCCA | 2746 |
| *RNA28SN5/1-5070* | 3872 | AGGGAACGGGCUUGGCGGAAUCAGCGGGGAAAGAAGACCCUGUUGAGCUUGACUCUAGUCUGGCAC | 3937 |
| *28SRNA:CR45844/1-3970* | 2747 | AGGGAACGGGCUUGGAAUAAUUAGCGGGGAAAGAAGACCCUUUUGAGCUUGACUCUAAUCUGGCAG | 2812 |
| *RNA28SN5/1-5070* | 3938 | GGUGAAGAGACAUGAGAGGUGUAGAAUAAGUGGGAGGCCCCCGGCGCCCCCCCGGUGUCCCCGCGA | 4003 |
| *28SRNA:CR45844/1-3970* | 2813 | UGUAAGGAGACAUAAGAGGUGUAGAAUAAGUGGGAGAUAUUAGACCU-----CGGU---------- | 2863 |

| | | | |
|---|---|---|---|
| *RNA28SN5/1-5070* | 4004 | GGGGCCCGGGGCGGGGUCCGCCGGCCCUGCGGGCCGCCGGUGAAAUACCACUACUCUGAUCGUUUU | 4069 |
| *28SRNA:CR45844/1-3970* | 2864 | ---------------------------UUGGUAUCGUCAAUGAAAUACCACUACUCUUAUUGUUUC | 2902 |
| *RNA28SN5/1-5070* | 4070 | UUCACUGACCCGGUGAGGCGGGGGGGC----------GAGCCCCGAG------------------- | 4106 |
| *28SRNA:CR45844/1-3970* | 2903 | CUUACUUACUUGAUUAAAUGGAACGUGUAUCAUUUCCUAGCCAUUAUACGGAUAUAUUUAUUAUAU | 2968 |
| *RNA28SN5/1-5070* | 4107 | -----G-----GGCU-------CUCGCUUCUGGCGCCAAGCGCCCGG------------------- | 4136 |
| *28SRNA:CR45844/1-3970* | 2969 | CUUAUGGUAUUGGGUUUUGAUGCAAGCUUCUUGAUCAAAGUAUCACGAGUUUGUUAUAUAAUCGCA | 3034 |
| *RNA28SN5/1-5070* | 4137 | ------------------------------------------------------C----CGCGCGC | 4144 |
| *28SRNA:CR45844/1-3970* | 3035 | AACAAAUUCUUUAAUAAAACGAUGCAUUUAUGUAUUUUUGAUUUGAAAAUUUGGUAUAACUCCAAU | 3100 |
| *RNA28SN5/1-5070* | 4145 | CGGCCGGGCGCGACCCGCUCCGGGGACAGUGCCAGGUGGGGAGUUUGACUGGGGCGGUACACCUGU | 4210 |
| *28SRNA:CR45844/1-3970* | 3101 | UACUCAGGUAUGAUCCAAUUCAAGGACAUUGCCAGGUAGGGAGUUUGACUGGGGCGGUACAUCUCU | 3166 |
| *RNA28SN5/1-5070* | 4211 | CAAACGGUAACGCAGGUGUCCUAAGGCGAGCUCAGGGAGGACAGAAACCUCCCGUGGAGCAGAAGG | 4276 |
| *28SRNA:CR45844/1-3970* | 3167 | CAAAUAAUAACGGAGGUGUCCCAAGGCCAGCUCAGUGCGGACAGAAACCACACAUAGAGCAAAAGG | 3232 |
| *RNA28SN5/1-5070* | 4277 | GCAAAAGCUCGCUUGAUCUUGAUUUUCAGUACGAAUACAGACCGUGAAAGCGGGGCCUCACGAUCC | 4342 |
| *28SRNA:CR45844/1-3970* | 3233 | GCAAAUGCUGACUUGAUCUCGGUGUUCAGUACACACAGGGACAGCAAAAGCUCGGCCUAUCGAUCC | 3298 |
| *RNA28SN5/1-5070* | 4343 | UUCUGACCUUUUGGGUUUUAAGCAGGAGGUGUCAGAAAAGUUACCACAGGGAUAACUGGCUUGUGG | 4408 |
| *28SRNA:CR45844/1-3970* | 3299 | UUUUGGUUUAAAGAGUUUUUAACAAGAGGUGUCAGAAAAGUUACCAUAGGGAUAACUGGCUUGUGG | 3364 |
| *RNA28SN5/1-5070* | 4409 | CGGCCAAGCGUUCAUAGCGACGUCGCUUUUUGAUCCUUCGAUGUCGGCUCUUCCUAUCAUUGUGAA | 4474 |
| *28SRNA:CR45844/1-3970* | 3365 | CGGCCAAGCGUUCAUAGCGACGUCGCUUUUUGAUCCUUCGAUGUCGGCUCUUCCUAUCAUUGUGAA | 3430 |
| *RNA28SN5/1-5070* | 4475 | GCAGAAUUCACCAAGCGUUGGAUUGUUCACCCACUAAUAGGGAACGUGAGCUGGGUUUAGACCGUC | 4540 |
| *28SRNA:CR45844/1-3970* | 3431 | GCAAAAUUCACCAAGCGUUGGAUUGUUCACCCAUG-CAAGGGAACGUGAGCUGGGUUUAGACCGUC | 3495 |
| *RNA28SN5/1-5070* | 4541 | GUGAGACAGGUUAGUUUUACCCUACUGAUGAU---GUGUUGUUGCCAUGGUAAUCCUGCUCAGUAC | 4603 |
| *28SRNA:CR45844/1-3970* | 3496 | GUGAGACAGGUUAGUUUUACCCUACUAAUGACAAAACGUUGUUGCGACAGCAUUCCUGCGUAGUAC | 3561 |
| *RNA28SN5/1-5070* | 4604 | GAGAGGAACCGCAGGUUCAGACAUUUGGUGUAUGUGCUUGGCUGAGGAGCCAAUGGGGCGAAGCUA | 4669 |
| *28SRNA:CR45844/1-3970* | 3562 | GAGAGGAACCGCAGGUACGGACCAAUGGCACA-AUACUUGUUCGAGCGAACAGUGGUAUGACGCUA | 3626 |
| *RNA28SN5/1-5070* | 4670 | CCAUCUGUGGGAUUAUGACUGAACGCCUCUAAGUCAGAAUCCCGCCCAGG-CGGAACGAUACGGCA | 4734 |
| *28SRNA:CR45844/1-3970* | 3627 | C-GUCCGUUGGAUUAUGCCUGAACGCCUCUAAGGUCGUAUCCGUGCUGGACUGCAAUGAUAAAUAA | 3691 |
| *RNA28SN5/1-5070* | 4735 | GCGCCGCGGAGCCUCGGUUGGCCUCGGAUAGCCGGUCCCCCGCCUGUCCCCGCCGGCGGGCCGCCC | 4800 |
| *28SRNA:CR45844/1-3970* | 3692 | GGGGCAA----------UUUGCAUUGUAUGGCUUCUAAACCAUUUAA---AGUUUA------UAAU | 3738 |
| *RNA28SN5/1-5070* | 4801 | CCCCCCUCCACGCGCCCCGCGCGCGCGGGAGGGCGCGUGCCCCGCCGCGCGCCGGGACCGGGGUCC | 4866 |
| *28SRNA:CR45844/1-3970* | 3739 | UUACUUUAUAAACGAC-------------AAU----------------------------GGAUGU | 3763 |
| *RNA28SN5/1-5070* | 4867 | GGUGCGGAGUGCCCUUCG--UCCUGGGAAACGGGGCGCGGCC-GGAGAGGCGGCCGCCCCCUCGCC | 4929 |
| *28SRNA:CR45844/1-3970* | 3764 | GAUGCC-AAUGUAAUUUGUAACAUAGUAAAUUGGGAGGAUCUUCGAU---CACCUGAUGCCGCGCU | 3825 |
| *RNA28SN5/1-5070* | 4930 | CGUCACGCACCGCACG-UUCGUGGG----GAACCUGGCGCUAAACCAUUCGUAGACGACCUGCUU | 4989 |
| *28SRNA:CR45844/1-3970* | 3826 | AGUUACAUAUAAAAGCAUUAUUUAAUACAAUGACAAAGCCUAGAAUCAAUUGUAAACGACUUUUGU | 3891 |
| *RNA28SN5/1-5070* | 4990 | CUGGGUCGGGGUUUCGUACGUAGCAGAGCAGCUCCCUCGCUGCGAUCUAUUGAAAGUCAGCCCUCG | 5055 |
| *28SRNA:CR45844/1-3970* | 3892 | AACAGGCAAGGUGUUGUAAGUGGUUGAGCAGCUGCCAUACUGCGAUCCACUGAAGCUUAUCCUUUG | 3957 |
| *RNA28SN5/1-5070* | 5056 | ACACAAGGGUUUGUC | 5070 |
| *28SRNA:CR45844/1-3970* | 3958 | CUUGAU-GAUUCGA- | 3970 |
